# Strong effects of temperature, population and age-at-maturity genotype on maturation probability for Atlantic salmon in a common garden setting

**DOI:** 10.1101/2022.07.22.501167

**Authors:** Eirik R. Åsheim, Paul V Debes, Andrew House, Petri T. Niemelä, Jukka P. Siren, Jaakko Erkinaro, Craig R Primmer

## Abstract

1. Age at maturity is a key life history trait and involves a trade-off between survival risk and reproductive investment, has close connections to fitness, and is an important factor for population structures. Temperature can have a dramatic influence on life history in ectotherms, but this influence may differ between populations. While an increasing number of studies have examined population-dependent reactions with temperature, few have investigated this in the context of maturation timing.
2. Atlantic salmon is a highly relevant study species for improving understanding of this topic as it displays considerable variation in life-history strategies, including maturation timing. Additionally, a large amount of this variation in maturation timing has been associated with a genomic region including the strong candidate gene *vgll3*, but the effect of this gene in the context of different environments and populations has not been studied.
3. Using a large-scale common-garden experiment, we find strong effects of temperature, population, and *vgll3* genotype on maturation in 2-year-old male Atlantic salmon. Observed maturation probability was 4.8 times higher in individuals reared at a mean temperature of 8.6°C compared to 6.9°C. This temperature effect was population-specific and was higher in the southern population compared to the northern population, potentially due to a higher intrinsic growth in the southern population as well as growth-temperature interaction.
4. The early-maturation *vgll3*\*E associated with a significantly higher maturation probability, but there was no *vgll3-*interaction with temperature or population.
5. Both body condition and body mass associated strongly with maturation; the body-condition association was stronger in fish carrying the *vgll3*\*E allele, and the body mass association was only present in the warm treatment.
6. Our findings demonstrate that the relative effect of *vgll3* on maturation timing is similar for two populations and two thermal environments and gives new perspectives on the relative effect of *vgll3* compared to such influences. Additionally, we show that populations can vary in their response to temperature change in terms of maturation timing, and that high intrinsic growth could potentially be associated with higher thermal sensitivity for life history variation.

## INTRODUCTION

Responses of wild animal populations to the changing climate are modulated by the phenotypic changes in individuals resulting from these changes in the environment. In this context, life history traits are of special interest as they describe the reproductive investment of organisms over their lifetime (Hutchings, 2021). Reaction norms describe the pattern of phenotypic expression of a genotype in differing environments and provide information about phenotypic plasticity and the presence of genotype x environment (GxE) interactions shaping the phenotype (Hutchings, 2011, 2021). The reaction norm between environment and life history may depend on the genetic background of the organisms, and thus animals from different populations-, or animals of different key genotypes, may respond differently to environmental influences (Oomen & Hutchings, 2015). This complicates the prediction and mitigation of climate change consequences for wild populations. Furthermore, teasing apart the contributions of genetics and environment can be challenging as these two factors are often correlated in wild populations. Rearing of individuals in common, controlled, conditions, i.e., common garden approaches, can partly resolve this issue by observing the phenotypic differences of animals with different genetic backgrounds reared in a common environment; By combining this approach with controlled variation of several environmental factors it is possible to build an understanding of the relative contributions of genes and environment to the phenotype, as well as the interactions between them.

Age at maturity is an important life-history trait as it describes at what age an organism will start reproducing (Cole, 1954). The Atlantic salmon (*Salmo salar*) is a highly relevant species for studying life history traits such as age at maturity in the context of better understanding genetic and environmental influences. Atlantic salmon displays considerable amount of variation in age at maturity (Reviewed in Mobley et al., 2021) arising from a combination of the number of years spent as a juvenile in freshwater and the number of years spent at sea before returning, often to their home river, to spawn. For example, in Atlantic salmon, the time spent at sea can vary from 0 to 5 years (Fleming, 1998; Fleming & Einum, 2011), with individuals typically doubling in size with each extra year spent at sea (Hutchings & Jones, 1998; Mobley et al., 2020). Further, some males never leave their home river and instead mature at a small size at the parr life stage, and so, mature individuals returning from the sea can be several thousand times larger than their mature river-bound counterparts. In recent decades, wild Atlantic salmon stocks have been in decline, with factors suggested to have contributed to this decline including climate change, aquaculture, illegal fishing, hydropower dams and harvesting of prey species (Chaput, 2012; Czorlich et al., 2022; Dadswell et al., 2021; Harvey et al., 2022; ICES, 2019; Vollset et al., 2022). Some of these factors have also been associated with life-history changes in the wild stocks, with some populations experiencing a decrease in the number or proportion of early-maturing individuals (Vollset et al., 2022), while others are reporting a decrease in large, late-maturing, individuals (Czorlich et al., 2018, 2022; Olmos et al., 2019). These trends thus make the study of factors impacting Atlantic salmon life-history traits highly timely.

Earlier, a locus including the gene *vgll3* was found to explain a large amount of variation (39%) in Atlantic salmon sea age at maturity in both males and females (Barson et al., 2015), a finding that has been replicated in both wild (Ayllon et al., 2015) as well as laboratory common garden studies (Debes et al., 2021; Sinclair-Waters et al., 2022; Verta et al., 2020). Changes in *vgll3* allele frequency have also been found to associate with a trend toward earlier maturation for Atlantic salmon in the river Teno (bordering Finland and Norway)(Czorlich et al., 2018). The two alleles of *vgll3* associate with either early (E) or late (L) maturation. While other studies have provided several clues for the developmental and molecular mechanisms involved in *vgll3’s* function (Debes et al., 2021; Kjærner-Semb et al., 2018; Kurko et al., 2020; Pashay Ahi et al., 2022; Verta et al., 2020), currently little is known about how this gene may interact with environmental factors like temperature and available nutrition, and so far, no common-garden studies have examined if there are population-differences in the effects of *vgll3* on maturation timing. Knowledge of how the effect of *vgll3* might vary between different environmental contexts and different Atlantic salmon populations is essential to understand how the effects of this gene on life history might change between populations and in the face of climate change.

Temperature can have a dramatic influence on the life history traits of ectotherms (Angilletta et al., 2004) and is known to have significant effects on maturation timing in Atlantic salmon (Friedland et al., 2009; Jonsson et al., 2016; Otero et al., 2011) and other ectotherm species (van der Have & de Jong, 1996). With the inherent variation in life history between Atlantic salmon populations, a key question is how this variation relates to changes in temperature, and how climate change impact might depend on the life-history strategy composition of a population. While there is a growing body of literature of studies on thermal reaction norms between fish of different populations and other genetic backgrounds, looking at traits like growth, survival, metabolism, and gene transcription (Hutchings, 2011; Oomen & Hutchings, 2015, 2022) few studies have investigated population differences in reaction norms between temperature and maturation timing, nor interactions with large-effect locus genotypes. Such research can help to better understand the potential impacts of global warming on the future life-history strategy composition of natural populations.

Here, we present the results of a common garden experiment investigating maturation timing in 2170 Atlantic salmon males with differing *vgll3* genotypes originating from two latitudinally distant populations from the Baltic Sea basin. Individuals were divided between a combination of two temperature treatments with a climate-change relevant 1.8°C temperature difference, and two feed treatments differing in nutrient proportions of lipids and caloric content. We aimed to explore the interactions between environment and genetics in shaping Atlantic salmon maturation timing, and to further broaden our understanding of *vgll3* as a large-effect life history gene in different genetic and environmental settings. More specifically, we test whether 1) the effect of *vgll3* genotype on maturation differs between the populations, temperatures, and feed treatments, 2) if the effects of temperature or feed treatment is population-dependent and 3) if other morphological phenotypes such as body mass or condition associate with maturation, and if *vgll3* genotype could influence this relationship.

## MATERIALS AND METHODS

### Study animals, crossing, initial rearing, and experimental feed and temperature treatments

The Atlantic salmon *(Salmo salar, Linnaeus 1758)* used in this study originated from two broodstocks – Neva and Oulu – maintained by the Natural Resources Institute Finland (LUKE). The Neva broodstock originates from the river Neva (Russia, 59.78°N, 30.71°E); It is maintained in a hatchery facility in Laukaa, Finland, and is regularly stocked in the river Kymijoki in southern Finland (60.48°N and 26.89°E). The Oulu broodstock originates 500 km further north, and was created as a mixture of several northern Baltic salmon populations including the river Oulujoki (Finland, 65.01°N, 25.27°E), where it is subsequently stocked regularly (Erkinaro et al., 2011; Karppinen et al., 2014). Both broodstocks are routinely renewed with individuals that complete marine migrations to their stocking rivers.

Broodstock and offspring individuals were genotyped using a multiplex-PCR for 177 single nucleotide polymorphisms (SNPs) of a previously described panel (Aykanat et al., 2016) as outlined in Debes et al. (2021). The panel included the *VGLL3*_TOP_ SNP (Barson et al., 2015) that was used for designing crosses to produce offspring with specific *vgll3* genotypes (see below). A subset of 131 SNPs in the panel not in high linkage disequilibrium was used for reconstructing the parents of the broodstock individuals as outlined in Debes et al. (2021) in order to avoid crossing closely related individuals.

Unrelated parents with homozygous *vgll3* genotypes were used to create a series of 2 × 2 factorials (one *vgll3*EE* male and female and one *vgll3*LL* male and female) so that each 2 × 2 factorial yielded four families, one of each of the four reciprocal *vgll3* genotypes (EE, EL, LE or LL), i.e., all offspring within a family had the same *vgll3* genotype (Suppl. Mat. Figure S1.1-Design). For analysis, we considered the two heterozygote combinations EL and LE as one genotype, EL. Only individuals from the same population were crossed together. In total, 13 and 17 2 × 2 factorials (52 and 68 families) were created using 50 and 67 parental individuals for the Oulu and Neva populations, respectively. Eggs and milt were stripped from the parental individuals at the broodstock hatcheries in mid (Oulu) or late (Neva) October 2017, immediately transported to the Viikki campus of the University of Helsinki, Finland, and fertilizations were conducted the following day.

The fertilized eggs of each family were divided between two temperature treatments (hereafter warm, cold), following a seasonal temperature cycle but with a 2°C difference maintained between the treatments (Figure 1). The eggs were incubated as outlined in Debes et al. (2021). Briefly, eggs of each family were randomly and equally divided between four separate flow-through incubators, two for each temperature treatment, i.e. two family replicates per temperature treatment, with families kept in separate compartments within an incubator (with randomized position). At first feeding, fish were transported to the University of Helsinki’s Lammi Biological Research Station (Lammi, Finland) and roughly equal numbers of individuals of each family from both populations were randomly chosen and placed into one of six replicate tanks of the same temperature treatment in which they had been incubated (2°C difference).

**Figure 1.**
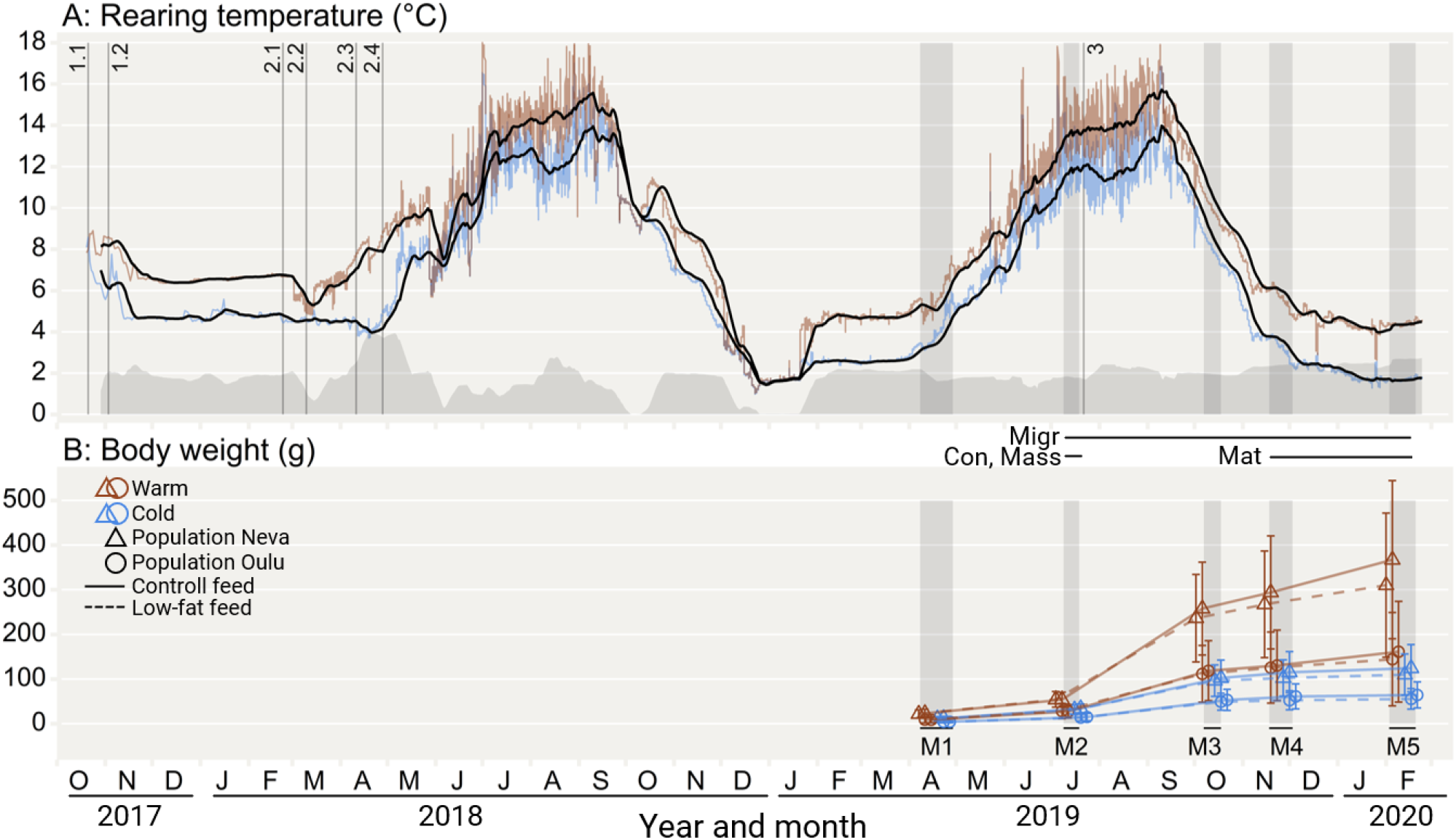
Timeline of the study showing temperatures (A) and fish growth (B). Shaded vertical areas (M1-5) indicate the timing of fish measurement sessions. The horizontal limes Con, Mass, Migr, and Mat indicate the measurement sessions from which data were taken to determine the body mass, condition, migration, and maturity status phenotypes, respectively, for use in modelling. A): Red and blue (upper and lower) lines indicate mean hourly water temperatures (of all tanks) for the warm and cold treatment, respectively. Black lines indicate the 10-day rolling water temperature average for each treatment. The grey area graph indicates the temperature difference between the 10-day rolling average of the two treatments, which averaged 1.8°C across the entire study. Periods with no temperature difference were due to heating/cooling system maintenance or technical malfunction due to very cold incoming lake water. Over the entire study, the mean temperature of the warm and cold treatment was 8.6 °C and 6.9°C, respectively. Vertical lines indicate the timing of the fertilisation of Oulu (1.1) and Neva (1.2) eggs; Transport of fish to Lammi Biological Station for Oulu-Warm (2.1), Neva-Warm (2.2), OuluCold (2.3), Neva-Cold (2.4) fish; and start of the feeding treatments (3). B) Points indicate mean body mass for fish in each population within each temperature-feed treatment. Points are repositioned horizontally within each measurement session to avoid overlap. Error bars indicate one standard deviation.

Some of the fish from some factorials were used in other experiments, and for this reason, the number of Oulu fish was around double the number of Neva fish. The tank transfers took place at four different time points, due to differences in the time of first feeding caused by the different incubation temperatures and the differing fertilization times for the two populations. Respectively, the transfer dates were 23.02.2018 and 11.04.2018 for warm- and cold treatment Oulu fish, and 10.03.2018 and 24.04.2018 for the warm- and cold-treatment Neva fish (Figure 1).

The feed treatments were started in July 2019 (summer in the second-year post-fertilization) and were combined with the temperature treatments. The feed treatments were either “control”, in which fish received regular Raisioaqua Baltic Blend aquaculture-grade feed (17-26% fat, 18.10-20.40 kj g^-1^ depending on pellet size), and “low-fat”, where the feed was replaced with a custom-made fat-reduced feed of the same brand, resulting in pellets of similar size and shape to the control feed, but with lower fat content (12-13% fat, 17.25 kj g^-1^)(see Supl. Mat. Table S1.1-Feed and S1.2-Nutrients for an overview of feed use and nutritional values). Thus, the 12 experimental tanks were divided into four treatment combinations, resulting in three tanks of each combination of temperature treatment (warm, cold) and feed treatment (control, low-fat). Tanks of different treatment combinations were evenly spread out in the research facility to minimize the occurrence of treatment-location correlations (Suppl. Mat. Figure S1.1-Design).

### Animal husbandry details

Following transport to Lammi Biological Station, fish were reared in experimental tanks (1.00 m tall, 2.77 m wide). The experimental tanks utilized a flow-through system of water that was pumped from a nearby lake (Pääjärvi) at approx. 12m depth. To reduce pathogen load, the incoming lake water was treated with UV light before entering the tanks. Water entered the tanks through a horizontal spray bar that created a circular flow in the tanks (which was standardized between tanks), and the water level and flow rate were increased over time as the fish grew. The water temperature in the tanks followed the seasonal lake temperature curve, while a heat-exchange system aimed to maintain a 2°C difference between the warm and cold treatments (Figure 1). Lighting was automated (on/off) and set to follow the local sunrise/sunset times (at 61.05°N, 25.04°E). Over the entire study, the mean temperatures of the warm and cold treatments were 8.6 °C and 6.9°C, respectively. The mean realized temperature difference was slightly lower (1.8°C) than the targeted 2°C difference due to heating/cooling system maintenance or technical malfunction due to very cold incoming lake water resulting in several short periods with no temperature difference between tanks during the first 15 months of the experiment (Figure 1)

Fish were fed *ad libitum* throughout daylight hours with body-size-matched pellets (Supl. Mat. Table S1.1-Feed, S1.2-Nutrients) of commercial fish feed (Hercules, Raisioaqua, Raisio, Finland). Feeding was conducted manually for the first 3-4 month (until mid-June 2018), after which, an automated feed delivery system was used (Arvo-Tek Oy, Finland). To adjust feed sizes and amounts in the early phase, a subsample of fish (120-300) was measured in July, August, October, and the end of November 2018. After that, feed amounts and sizes were adjusted based on size data from the regular phenotypic measurements which started in April 2019 (described below). Internal tank surfaces were scrubbed clean at least once per week. Tanks were visually inspected on a daily basis; dead fish were removed from the tanks, and any moribund fish were removed and euthanized with an MS222 overdose (0.250 g L^-1^, sodium bicarbonate-buffered). To provide the fish with environmental enrichment, half of each tank was covered with a dark-green camouflage mesh. These covers were installed at the end of April 2018 for the warm tanks, and in the middle of June 2018 for the cold tanks so that fish had experienced similar degree-days when the nets were installed.

### Weighing, measurements and maturation checks

At the first measurement in April 2019 (Figure 1; M1), all individuals were tagged with a passive integrated transponder (PIT-tag) inserted into the abdominal cavity about half a centimetre caudally from the right-side pectoral fin using sterilised needles. At the same time, a small fin clip was taken from their caudal fin, allowing for genotyping, sex determination, and parental assignment as in Debes et al. (2021), and thereby, individual identification from that point on.

Individual phenotypic characteristics were recorded five times between April 2019 and February 2020 at 2– 3-month intervals (Figure 1; M1-M5). These include body mass, body length, and two life-history phenotypes. These phenotypes were migration phenotype (smolt or parr; which in wild fish indicates the initiation of marine migration) and status of sexual maturity. Migration phenotype was checked at every measurement session from June 2019 to February 2020 (M2-M5). Maturation status was checked in Nov/Dec 2019 and February 2020 (M4-M5). Maturation status was checked by carefully stroking each individual’s abdomen towards the vent; Fish releasing milt were categorised as mature. No females showed any signs of maturation e.g., bloated belly. Migrant (smolt) vs. resident (parr) phenotype was checked using criteria including level of silvering and occurrence of parr marks. Individuals were recorded as having smolted from the time point following the last recording of resident (parr) characteristics.

For measurements, fish were netted from their holding tank to a continuously aerated anaesthetic bath (MS222, 0.125 g L^-1^, sodium bicarbonate-buffered) at a similar temperature (within 1°C) to the tank water. Each individual’s body mass was then recorded to the nearest 0.01 g (April, July) and subsequently 0.1 g using a digital scale (Scout STX222 or STX6201, Ohaus, Parsippany, USA). Fork length (length from snout to fork of tail) was measured to the nearest mm using a digital fish-measurement board (DCS5, Big Fin Scientific, Austin, TX, USA), after which migration and maturation phenotypes were recorded and the fish were returned to its tank. Those performing the measurements were blind to the genotype and population of origin of the fish, but not temperature and feeding treatment.

## STATISTICAL ANALYSIS

### Sample size

As no females matured during the focal period, this study focuses solely on males. Due to the initially unknown rates of early maturation and mortality, we aimed for a sample size as high as possible given our supply of eggs from the hatcheries. This was to ensure we would have sufficient statistical power to test for the direct- and interaction-effects of our genetic and environmental factors. By April 2019, a total of 2657 males were tagged. By the following winter (Feb 2020), 263 males died prematurely, while 124 had been euthanized for use in another project (balanced among tanks, sex, *vgll3* genotypes, and families). A further 98 males were excluded due to incomplete genotype data, and 2 were excluded due to other incomplete data. A total of 2170 males were thus included in the final analysis.

### Dataset and included variables

Each male individual counted as one observation. We included only fish with successfully identified *vgll3* genotypes, parental identities, and sex. The variables included were *vgll3* genotype *(EE, EL, LL)*, population of origin (Neva, Oulu), feeding treatment (control, low-fat), temperature treatment (cold, warm), migration phenotype status as observed by February 2020 (migrant, resident), maturation status by February 2020 (matured, not matured), log body mass (g, mean centred and SD scaled) and body condition (%, mean centred and SD scaled) in July 2019. This timepoint for body condition and mass was chosen as the one most likely relevant for future maturation, representing the state of body reserves before the enlarging gonads start influencing body condition (Rowe et al., 1991). Body condition was calculated as the residuals of a linear model of the log body mass (g) against the log body length (mm) on the entire study population, thus being represented as percent difference in body mass from the expected body mass (given length). *vgll3* genotype was split into two variables, one for the gene’s additive effect (*vgll3*_*add*_, coded *EE*=1, *EL*=0, *LL*=-1) and one for the dominance-effect, i.e., deviance from an additive pattern (*vgll3*_*dom*_, coded *EE*=0, *EL*=1, *LL*=0) as in Xiang et al. (2018).

### Maturation probability models

We used a general linear mixed-effect modelling approach (Bernoulli distributed, logit link) to examine how maturation probability (response variable) associated with the explanatory variables *vgll3* genotype, population of origin, feed treatment, temperature treatment, body condition, body mass and migration phenotype. Although the covariates body condition, body mass, and migration phenotype were not experimentally manipulated variables, they were included in the full model as explanatory variables to improve the model’s overall fit and to examine how these biologically relevant covariates interact with the genetic and environmental factors (Model-Mat-Cov or “full model”). For comparison, we also fitted an alternative no-covariate model which excluded body condition, body mass, and migration phenotype (Model-Mat-Nocov or “no-covariate model”). We fitted interactions between *vgll3* and temperature treatment, population of origin, feeding treatment, body condition, body mass, and migration phenotype. Additionally, we fitted an interaction between population and temperature, as well as temperature- and population interactions with the covariates body condition, body mass, and migration phenotype. Rearing tanks were included as random effects (on the intercept) to account for between-tank (i.e., environmental) variation. Relatedness was accounted for by including the pedigree information (up to the grandparents) into the model using an animal model approach (Henderson, 1973; Wilson et al., 2010), i.e., using the inverse of the additive genetic relatedness matrix to fit an effect of the individual animal as a random effect (on the intercept), which also gives an estimate of the additive genetic variance. Heritability was calculated using the no-covariate model only, using the estimates of additive genetic variance as described in de Villemereuil (2021). Variance explained by *vgll3* was estimated as in Debes et al. (2021).

### Supplemental models

Four supplemental models were fitted to explore whether *vgll3* associated with any of the three non-independent covariates as response variables: body mass (Model-Mass), body condition (Model-Cond), and migration phenotype (Model-MigPheno). These models were fitted using the same explanatory variables as the no-covariate maturation probability model (*vgll3*, temperature, population, feed), but with the following differences: Model-Mass and Model-Cond were fitted using an identity-link instead of a logit link, making them linear mixed-effect models instead of generalised linear mixed-effect models; in addition, Model-Mass and Model-Cond did not include feeding treatment as an explanatory variable since the measure of body mass and condition used in these models was recorded before the feeding treatment started (Figure 1). Finally, to allow for a closer examination of *vgll3*’s effect on body condition, an additional model was fitted for body condition including migration phenotype as a covariate (Model-Cond-Cov).

### Model fitting approach

All models were fitted and analysed using a Bayesian approach for generalised- and non-generalized linear mixed models. Models were fitted using *Rstan* via the *R* package *brms*. All models were fitted using 4 MCMC chains run for 3000 transitions, discarding the 500 first transitions of each chain for warmup, thus totalling 10000 posterior samples for each model. For the maturation- and migration phenotype models, prior distributions of the intercept, effect sizes, and the SDs of the random effects were all set to a relatively non-informative normal distribution with a mean of 0 and a standard deviation of 2. For the body condition and body mass models, the same parameters were given priors with a normal distribution of 0 and a standard deviation of 1. All model fits were verified using a visual posterior predictive check and checked for influential points by inspecting pareto k diagnostic values. Model-Mass had a large proportion of highly influential points (23.1% of values with K > 0.7), motivating a more careful interpretation of this model. Full model summaries can be found in supplementary material S2. Interactions were generally considered non-significant when the 95% credible interval of their effect size included 0. For some of those cases (noted in results), we simplified the model estimates by calculating the unconditional (marginal) mean estimates of the main effects. Unconditional estimates were calculated as the mean of one effect (i.e. *vgll3*) over all levels of the non-significant interaction variable (i.e feeding treatment). All these calculations were done using the posterior distributions of the parameter estimates (effect sizes) taken from the rstan output.

### Statistical software

All analyses were performed in the *Rstudio* v.2022.02.3 (RStudio Team, 2022) software environment running *R* v.4.1.2 (R Core Team, 2021) and *Rstan* v2.21.5 (Stan Development Team, 2022). R packages used for analysis were *brms* v.2.17.0 (Bürkner, 2017, 2018, 2021) for working with *Rstan* models, *loo* v.2.5.1 (Vehtari et al., 2017, 2022) for inspecting pareto k diagnostic values, *ggplot2* v3.3.5 (Wickham et al., 2021) for visualization, and *tidyverse* v1.3.1 (Wickham et al., 2019) for various programming and data management tasks.

## RESULTS

### Observed maturation rates

The overall male maturation rates in the Oulu and Neva populations were 13.3% and 32.6%, respectively (of n=1335 & 835). Across-population maturation rates in the cold and warm treatments were 6.7% and 36.6% (of n=1154 & 1016), while *vgll3* genotype-specific maturation rates were 6.6%, 18.2%, and 35.6% for *vgll3* genotypes *LL, EL*, and *EE* (of n=457, 1095 & 618), respectively (Table 1).

**Table 1.** Observed maturation rates for male Atlantic salmon for all combinations of *vgll3* genotype, temperature treatments and population of origin. The numbers in parentheses indicate the total number of fish in that group.

| Group | LL | EL | EE | Total |
| --- | --- | --- | --- | --- |
| Neva | 13.6% (147) | 28.7% (422) | 49.2% (266) | 32.6% (835) |
| Oulu | 3.2% (310) | 11.6% (673) | 25.3% (352) | 13.3% (1335) |
| Warm | 13.0% (215) | 32.9% (517) | 61.3% (284) | 36.6% (1016) |
| Cold | 0.8% (242) | 5.0% (578) | 13.8% (334) | 6.7% (1154) |
| Neva, warm | 27.4% (73) | 48.8% (217) | 76.4% (140) | 54.2% (430) |
| Neva, cold | 0.0% (74) | 7.3% (205) | 19.0% (126) | 9.6% (405) |
| Oulu, warm | 5.6% (142) | 21.3% (300) | 46.5% (144) | 23.7% (586) |
| Oulu, cold | 1.2% (168) | 3.8% (373) | 10.6% (208) | 5.1% (749) |
| Total | 6.6% (457) | 18.2% (1095) | 35.6% (618) | 20.7% (2170) |

### Vgll3

Maturation probability was higher for each carried *vgll3*\*E allele (Table 2, Figure 2). This increase was largely additive, as we found no significant dominance-effect of either allele. Only in the full maturation model were there indications of a dominance effect for the E allele, but the effect was small and its 95% credible interval overlapped with zero (Model-Mat-Cov, Figure 3). The only variable having a clearly significant interaction with *vgll3* was body condition, and as such, no significant interactions with *vgll3* (on maturation probability) were found for population, temperature, feed, body mass, or migration phenotype. The full maturation model (Model-Mat-Cov, Figure 3) estimated that each carried *vgll3*\*E allele increased the odds of maturation 7.42-fold [95% CI: 3.23, 20.17] (unconditional on interactions with feed, temperature, population, and migration phenotype), and that each carried E allele was estimated to increase the body-condition effect 1.56-fold [95% CI: 1.13, 2.21]. *Vgll3*-effects on the migration phenotype, body condition, and body mass covariates were all close to zero (Suppl. Mat. Figure S2.2, S2.3, S2.4).

**Table 2.** Predicted maturation probabilities based on the no-covariate maturation model (Model-Mat-Nocov) for male Atlantic salmon in different grouped combinations of *vgll3* genotype, temperature treatment and population of origin. Brackets indicate 95% credible intervals. For these predictions, feed treatment was set to control-feed.

| Group | LL | EL | EE |
| --- | --- | --- | --- |
| Neva-Cold | 0.2% [0.0%,1.5%] | 1.3% [0.2%,5.1%] | 7.2% [1.2%,26.3%] |
| Neva-warm | 14.0% [2.2%,50.6%] | 61.6% [30.7%,86.7%] | 93.2% [75.6%,98.8%] |
| Oulu-Cold | 0.1% [0.0%,0.8%] | 0.6% [0.1%,2.0%] | 2.6% [0.5%,9.5%] |
| Oulu-warm | 1.6% [0.2%,7.1%] | 10.7% [3.5%,24.4%] | 43.4% [16.9%,73.9%] |

**Figure 2.**
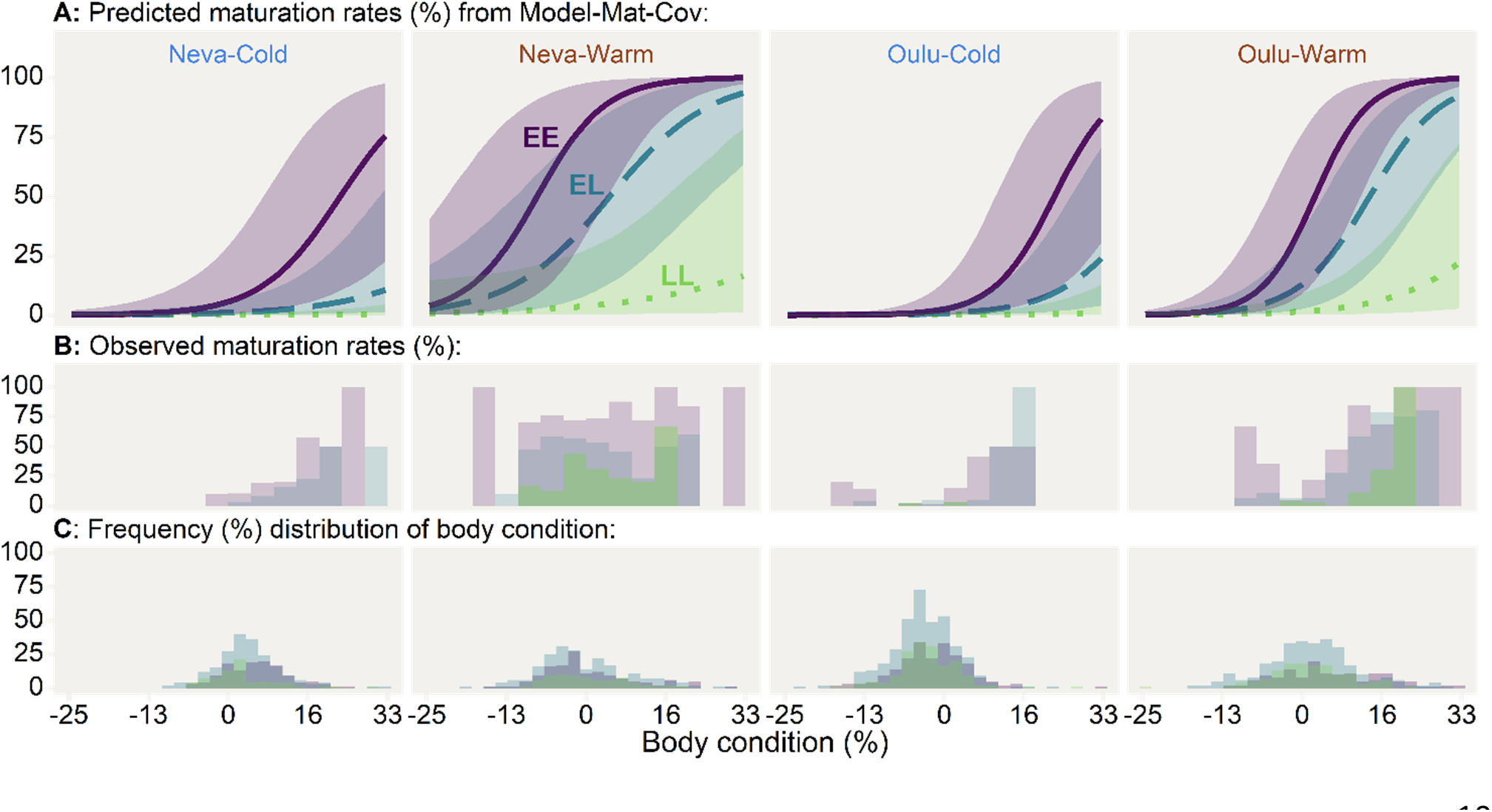
Predicted (A) maturation probabilities and observed (B) maturation rates for three different *vgll3* genotypes (Purple-solid=EE, Blue-dashed=EL, Green-dotted=LL), two temperature treatments (Cold, Warm), and two populations of origin (Neva, Oulu), plotted against body condition. C) shows corresponding frequency distributions of body condition. Body condition represents the percentage difference in body mass from the expected body mass given body length. Lines represent the mean predicted maturation probability, with shaded areas around the lines indicating the 95% credible interval for the predictions (overlap is not an indicator of significance). Predictions are based on the full model for maturation probability (Model-Mat-Cov, Figure 3, Table S2.4). Body mass for predictions is set to the mean of the whole study population and the migration-phenotype parameter is set to 0.5 (giving an estimate unconditional on migration-phenotype).

**Figure 3.**
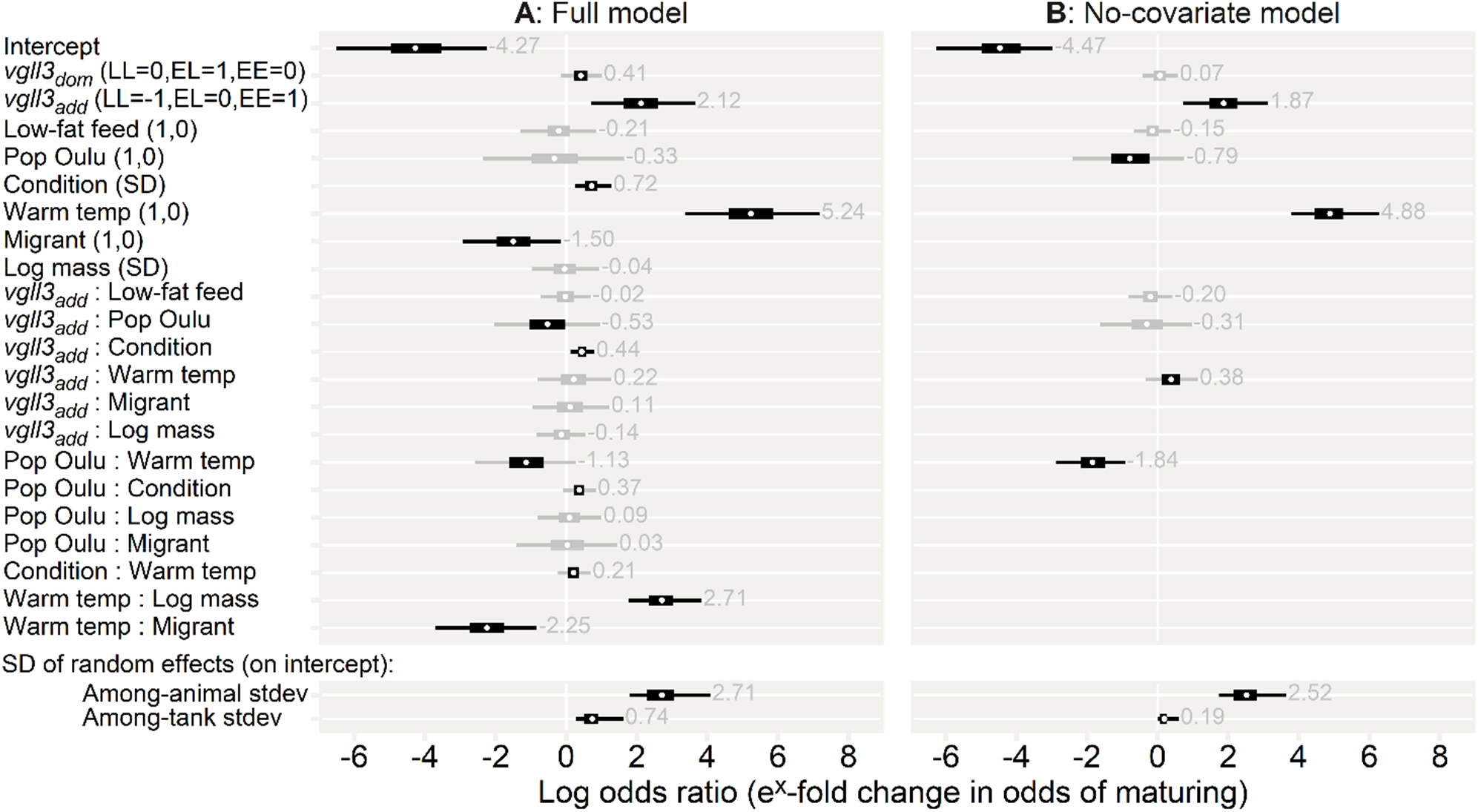
Effect sizes (parameter estimates) for the two maturation probability models. A) Parameter estimates for the full model (Model-Mat-Cov). B) A simplified model that only includes independent variables (Model-Mat-Nocov). Effects are shown as log odds ratios, thus indicating relative change in odds (odds = probability of maturing divided by the probability of not maturing); Odds 0, 1 and infinite convert to probabilities 0, 50% and 100%, respectively. Thick and thin sections of bars indicate 97.5% and 50% credible intervals, respectively. As a visual aid, intervals are coloured grey if they include 0. Grey numbers show the mean parameter estimate. Parentheses indicate the levels of the variables, and all variables are set to 0 for the intercept. The first level in parenthesis is the written level. Body condition and log body mass are SD scaled and mean-centred (mean=0), so the parameter estimates indicate the effect of increasing or decreasing either of these variables with one SD. The *vgll3dom* parameter indicates the degree of dominance displayed by either of the alleles. The lower section shows the standard deviation of the random effects, representing the degree of among-tank variation and among-animal variation (additive genetic standard deviation). The full model summaries can be found in the supplementary materials (Tables S2.4, S2.5)

### Temperature and population

Maturation probability was higher in the warm temperature treatment compared to the cold, and temperature and population interacted so that the maturation probability difference between temperatures was higher in the Neva population (Figure 4, Figure 3, Table 2). Compared to the cold-treatment fish, the warm-treatment Neva and Oulu fish were, respectively, estimated to have a 131.87-fold [95% CI: 44.06, 539.21] and 20.89-fold [9.14, 56.71] higher odds of maturing (Model-Mat-Nocov, Figure 3). Maturation probability was thus higher for Neva fish, but only significantly so in the warm temperature treatment. Compared to Oulu fish, Neva fish had a 13.88-fold [95% CI: 3.08, 76.89] higher odds of maturing in the warm treatment, but only a 2.20-fold [95% CI: 0.47, 11.07] higher odds of maturing in the cold treatment (in the no-covariate model).

**Figure 4.**
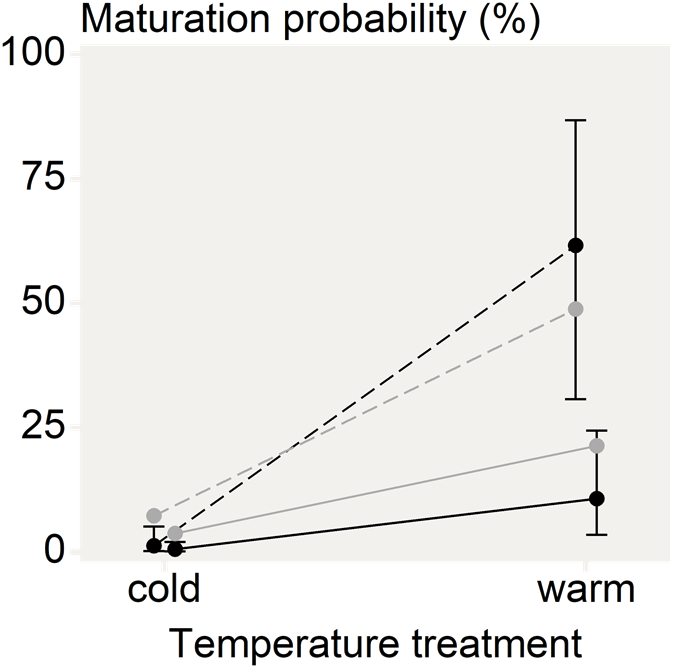
Predicted maturation probabilities (black) and observed maturation rates (grey) for fish from the Oulu (solid line) and Neva (dashed line) populations in each temperature treatment (mean temperatures: cold=6.9°C, warm=8.6°C) with the *vgll3*EL* genotype in the control-feed treatment. The predictions are based on the no-covariate maturation model (Model-Mat-Nocov, Figure 3, Table S2. 5). Error bars show the 95% credible interval of each prediction (overlap is not an indicator of significance).

The estimated population-effect and interaction with temperature was reduced in the full model which included the covariates body mass, body condition and migration phenotype (Model-Mat-Cov, Figure 3); In that model (using the same comparison as above), compared to Oulu fish, Neva fish had a 4.32-fold [95% CI: 0.54, 37.30] higher odds of maturing in the warm treatment, and a 1.39-fold [95% CI: 0.19, 10.48] higher odds of maturing in the cold treatment.

Body mass was higher in the Neva population and in the warm temperature treatment, with the modelled body mass of Neva fish estimated to be 151.44% [95% CI: 95.66, 223.48] higher than Oulu fish, and the body mass of fish in the warm treatment being 73.73% [95% CI: 44.85, 107.00] higher than in the cold treatment (Model-Mass, Suppl. Mat. Figure S2.3). The probability of smolting (transitioning to the migrant phenotype) was higher for the Neva population and in the warm treatment, with the Neva fish having an estimated 18.69-fold [95% CI: 3.95, 104.74] increase in the odds of smolting compared to Oulu, and the warm-treatment fish having a 20.87-fold [95% CI: 5.99, 93.13] increase in the odds of smolting compared to the cold-treatment fish (Model-MigPheno, Suppl. Mat. Figure S2.4). There were no significant interactions between population and temperature in their effect on body mass or migration phenotype.

### Body condition

Maturation probability increased with higher body condition (Figure 2), and this effect had a small and slightly uncertain interaction with population so that the effect of body condition was slightly higher for Oulu fish (Model-Mat-Cov, Figure 3). The effect of body condition was similar in both temperatures (no significant interaction). For an Oulu fish, an increase in body condition of one standard deviation resulted in a 3.30-fold [95% CI: 2.37, 5.07] increase in the predicted odds of maturing, while for a Neva fish, the increase in predicted odds was 2.28-fold [95% CI: 1.53, 3.71] (both estimates unconditional on temperature).

### Body mass

Higher body mass was associated with higher maturation probability, but only in the warm temperature treatment. The effect of body mass was similar in both populations (no significant interaction). The full maturation model (Model-Mat-Cov, figure 3) estimated that an increase in body mass by one SD would increase the odds of maturing 15.06-fold [95% CI: 7.03, 40.89] in the warm treatment and 1.00-fold [95% CI: 0.46, 2.27] in the cold treatment (both estimates unconditional on population).

### Migration phenotype

Observed maturation-rates were higher among fish that had smolted (transitioned to the migrant phenotype) prior to spawning, being 23% amongst migrants (smolt, n=1586) and 13% amongst residents (parr n=584). However, after accounting for body mass and body condition, smolting prior to the spawning season was associated with a lower maturation probability, and this effect was stronger in the warm temperature treatment. The effect of migration phenotype was the same for all *vgll3* genotypes and in both populations (no significant interactions). The full maturation model (Model-Mat-Cov, Figure 3) estimated that the migrant phenotype in the cold treatment had a 0.22-fold [95% CI: 0.05, 0.86] lower odds of maturing compared to the resident phenotype, and 0.02-fold [95% CI: 0.00, 0.11] lower odds in the warm temperature treatment (both estimates unconditional on population). 73% (n=1526) of the fish in this study had smolted by early 2020 (winter of third year post-fertilization), of which 30% (n=357) also matured. See supplementary Table S2.10 for an overview of combined smolting and maturation rates.

### Feed

The low-fat feed had no significant effect on maturation probability nor any detectable interactions with any other variables (Model-Mat-Cov, Model-Mat-Nocov, Figure 3).

### Random effects and heritability

In the full maturation model (Model-Mat-Cov, Figure 3), the estimated among-tank variation in maturation probability equated to an average 2.10-fold [95% CI: 1.32, 5.04] deviation from the model’s intercept, which indicates relatively minor tank-related environmental effects on maturation. The among-animal variation was larger, with an average deviation of the model’s intercept equating to a 15.04-fold [95% CI: 6.03, 59.59] change in odds of maturation from the mean. For example, for a mean maturation probability of 50%, the among animal (additive genetic) standard deviation would equate to a range in probabilities going from 91% to 10%.

Total heritability (estimated from the no-covariate maturation model) including variation caused by *vgll3* was estimated at 0.69 [95% CI:0.54,0.82] indicating that around 69% of the remaining variation in maturation probability after accounting for the other model terms (temperature treatment, feed, population) could be ascribed to additive genetic effects, suggesting there is a proportionally large amount of additive genetic variance for male maturation. The additive variance contributed by *vgll3* was estimated at 0.13 [95% CI:0.05,0.23].

## DISCUSSION

### Population and temperature

We found variation in population-level thermal reaction norms for early male maturation. Maturation probability was higher in the Neva population, but only significantly so in the warm temperature treatment. Both populations displayed considerably higher maturation rates in the warm-compared to the cold treatment, and this response was stronger for the Neva population. This could indicate higher thermal sensitivity of fish from the Neva population, and that future temperature increases as a consequence of climate change could have a stronger effect on this population in terms of maturation timing.

Temperature is known to have a large influence on maturation timing in salmon, although experimental studies have found mixed results regarding the direction of this effect. Some studies have observed an increase in parr maturation with increased temperature (Fjelldal et al., 2011; Imsland et al., 2014), some have observed reductions (Herbinger & Friars, 1992), and some have observed no change at all (Baum et al., 2005). Some of these discrepancies are likely due to differences in the temperatures used and the timing of the temperature treatments. Our findings show that a life-long chronic difference in mean temperature of 1.8°C (from 6.9 to 8.6°C) can cause a large increase in the probability of early maturation for male Atlantic salmon, in our case going from an observed maturation rate of 6.7% to 36.6%. Further, given the relatively high heritability of maturation probability at 0.7 (of which the *vgll3* locus contributes 0.13), we find that there is strong potential for natural selection on this trait, which could either counteract or exacerbate the influence of climate change on maturation timing, depending on its fitness effects.

Adaptation to colder environments sometimes involves an increased growth rate to compensate for the slowed-down growth and metabolism that happens at lower temperatures (countergradient variation). This has been found for several species of fish (Baumann & Conover, 2011; Conover et al., 1997; Conover & Present, 1990; Yamahira et al., 2007; Yamahira & Conover, 2002), but the pattern is not universal (Belk et al., 2005; Hutchings et al., 2007; Oomen & Hutchings, 2016). With countergradient variation, we would have expected the Oulu fish to display an inherently higher growth rate compared to the Neva fish, as the Oulu population originates about 500 km further north than the Neva population. Instead, we found that the Neva fish outperformed the Oulu fish in terms of growth at both temperatures (mean 8.6°C and 6.89°C). Nevertheless, the population-based difference in maturation was non-significant at the cold temperature, which could be interpreted as growth being less inhibited by the cold temperature treatment in Oulu fish. Interesting follow-up research could include assessment of whether the Neva fish would maintain higher growth rates than the Oulu fish at even lower temperatures, or if the Oulu fish would be more robust to a further temperature decrease. Another source of variation between the populations could be differences between the hatcheries where the broodstock fish are initially reared. However, we do not expect this to have a large impact as the hatcheries are not applying any specific selection to the fish they are using, and all broodstock fish have completed a marine migration before being taken into the facilities.

### *Vgll3* effects

In the time since the first finding of the association between *vgll3* and Atlantic salmon age at maturity (Ayllon et al., 2015; Barson et al., 2015), this effect of *vgll3* on maturation has been validated in three common-garden studies using male Atlantic salmon in their first-year post fertilization (Debes et al., 2021; Sinclair-Waters et al., 2022; Verta et al., 2020). The current study builds on these findings by showing that the effect of *vgll3* genotype on maturation timing also holds for male Atlantic salmon reared in common, but cooler and less controlled, thermal conditions than the previous studies, as well as showing that the relative effect of *vgll3* remains similar independent of a 1.8°C temperature difference, and independent of any potential genetic or epigenetic influences of the population of origin. Additionally, the design of this experiment allowed us to quantify the relative effect of *vgll3* on maturation compared to the effect of temperature, and we found that the effect of a single *vgll3*E* allele on maturation was 39% that of a 1.8°C temperature increase

We modelled maturation probability using a threshold model (logit-link) in which we assume that maturation (a binary trait) is determined by some underlying liability trait that must reach a certain threshold to initiate the maturation process. The positive effect of *vgll3* genotype and temperature on the liability of maturation can be interpreted as these predictors either 1) having a positive effect on the unknown liability trait itself, or 2) lowering the liability trait’s threshold to induce maturation. Somatic growth is often assumed to be a key liability trait for maturation (Taranger et al., 2010), however, by separately modelling body condition and body mass as response variables, we found *vgll3* to not significantly affect either of these traits directly, which means either that *vgll3* influence arises through other non-measured liability traits which do not perfectly associate with body condition or growth, or that *vgll3* works by changing the threshold of maturation for body condition and growth. This contrasts slightly with what was found in Debes et al (2021), where a small effect of *vgll3* was found on body condition. However, that study had higher sample sizes (N=2608) of both sexes from a single population (Neva) and temperature, used fish in their first-year post-fertilization (i.e. no individuals had undergone smoltification), and covered multiple timepoints reaching well into autumn. It was also noted that the presence and magnitude of the association varied across time, and between feeding treatments and sexes. Here, we used fish in their second-year post-fertilization and only included body condition in early July, so the different results may be attributed to any of these differing factors. Further, 73% of the fish in our study also underwent the smolting process (Table S2.10), which we found to affect body condition, and it is possible that an imperfect accounting of this effect could mask a *vgll3*-body condition association, or the smoltification process itself may modify the association.

The interaction between *vgll3* and body condition indicates an effect on the body condition threshold for affecting maturation, which becomes lower and narrower for each *vgll3*E* allele in the genotype (Figure 3A). On the other hand, the lack of an interaction between *vgll3* and temperature indicates that their relative effects (relative to the contribution of the other predictors) are independent of each other. The effect of *vgll3* was also similar between the two populations, indicating that any genetic or epigenetic factors that were different between the two populations did not significantly interact with the mechanisms of this large-effect locus.

We found no signs of a dominance pattern for the effect of *vgll3* on maturation probability, which is in line with what has been found in some other common-garden studies on early maturation in male Atlantic salmon (Debes et al., 2021; Sinclair-Waters et al., 2022), but not all (Verta et al., 2020) and not with the initial GWAS study on wild-caught individuals which found a sex-dependent dominance pattern for *vgll3* (Barson et al., 2015).

### Feed, body mass and condition

Acquisition of sufficient energy stores is a key part of the process leading up to sexual maturation (Berglund, 1992; Norrgård et al., 2014; Rowe et al., 1991; Reviewed in Mobley et al., 2021). In line with this, we found body mass and body condition to strongly affect maturation probability, with individuals having larger body mass or body condition being associated with an increased probability of maturation. The observation that both body condition and mass were estimated to have significant effects within the same model could indicate that these traits have independent effects on maturation timing, and that both traits need to be considered together to gain a complete understanding of the maturation processes. In terms of the relative effects of these traits, body mass had the strongest effect on maturation probability, and one standard deviation of log body mass had a 4.4 times larger effect than one standard deviation of body condition (in the warm temperature treatment, see below). Body condition is known to correlate with body fat content in Atlantic salmon (Herbinger & Friars, 1991) and other fish (Chellappa et al., 1995; Mozsár et al., 2015), and has been shown to be important for early male maturation in Atlantic salmon (Rowe et al., 1991; Simpson, 1992) and in chinook salmon (*Oncorhynchus tshawytscha)* (Shearer & Swanson, 2000). The observed effect of body condition on maturation is then likely due to the influence of lipid stores (Parker & Cheung, 2020), while body mass might reflect other unrelated growth- or development-related factors.

Similar to what has been previously demonstrated in nine-spined stickleback (Kuparinen et al., 2011), we found temperature to affect maturation not only through growth in body size but also through some other unknown pathway (as temperature still had a large effect on maturation after accounting for growth), further indicating that there are other biological processes important for maturation that do not involve body mass or body condition.

The effect of body mass on maturation was only observed in the warm temperature treatment, further emphasising the differences in the influences of body mass and body condition; In the cold treatment, no effect of body mass was found, yet the effect of body condition remained. This could indicate that body mass does not start to influence maturation before the individual has reached a certain developmental threshold which is affected by temperature. As far as we are aware, such an interaction between temperature and body mass has not been described before. For future research, it might be helpful to see this finding replicated using a higher number of temperature treatments and with smaller temperature increments. If this is a general pattern between temperature, growth and maturation, it could mean that the sensitivity of different populations to climate change in terms of life history could be closely connected to their somatic growth rate.

No effect of the feeding treatment was detected for the probability of maturation, so fish that were fed the fat-reduced diet from July and onwards i.e., four months prior to spawning, had the same probability of maturing as those fed the control feed. This suggests that either the reduction in fat was not sufficient, and/or the maturation process had been initiated prior to the start of the treatment. This is in line with the findings reported by Debes et al. (2021) who applied a 2-day vs. 7-day per week *ad libitum* feeding restriction treatment for a six-week period starting in September (2-3 months prior to spawning), but did not find any difference in maturation probability between the treatments despite large effects of the feeding treatment on growth and condition.

## CONCLUSION

Temperature, population of origin, and *vgll3* genotype each had a significant influence on maturation in two year old Atlantic salmon males. We found a population-dependent thermal reaction norm for maturation probability, suggesting that the two populations might respond differently to climate change in terms of life-history strategies. We also found the effect of body mass on maturation to be highly temperature-dependent, which suggests that responses to temperature changes in different populations could be connected to individual growth rates within each population. Further understanding of this pattern could benefit by exploring a larger range of temperatures. The relative influence of *vgll3* on maturation probability was the same for both temperatures and populations, suggesting that the relative contribution of *vgll3* to maturation timing is similar between Atlantic salmon populations for early male maturation. In terms of the mechanisms by which *vgll3* influences maturation, we did not find significant influences of *vgll3* on body mass or body condition, suggesting that a significant portion of *vgll3’s* influence is coming from pathways other than growth, or through lowering the growth threshold of maturation.

## ACKNOWLEDGEMENTS

We acknowledge Suvi Ikonen, Nikolai Piavchenko, Paul Bangura, Tiemen Jansen, Shadi Jansouz, Teemu Mäkinen, Jack O’ Callagan, Anna Toikkanen, Ilke van Gestel, and Pirta Palola for their assistance with animal husbandry and with data collection; Jacqueline Moustakas-Verho, Kzenia Zueva, Marion Sinclair-Waters, Nico Lorenzen, Victoria Pritchard, Yann Czorlich, Suvi Ikonen, Spiros Papakostas, and the Laukaa and Taivalkoski hatchery staff are thanked for assistance with initial sperm- and egg collection and/or fertilizations; Susanna Airaksinen and Thomas Ginström and others at Raisioaqua Ltd for producing the control- and low-fat feeds; Tutku Aykanat for calling SNPs; Annukka Ruokolainen, Seija Tillanen, and Shadi Jansouz for *vgll3* genotyping and sexing; Lammi Biological Station for hosting our common garden facilities and especially John Loehr for assistance with local logistics; CSC – IT Center for Science, Finland, for computational resources; and Natural Resources Institute Finland (LUKE) for access to the broodstocks.

## SUPPLEMENTARY MATERIAL 1: Supplementary methods and design

**Figure S1.1.**
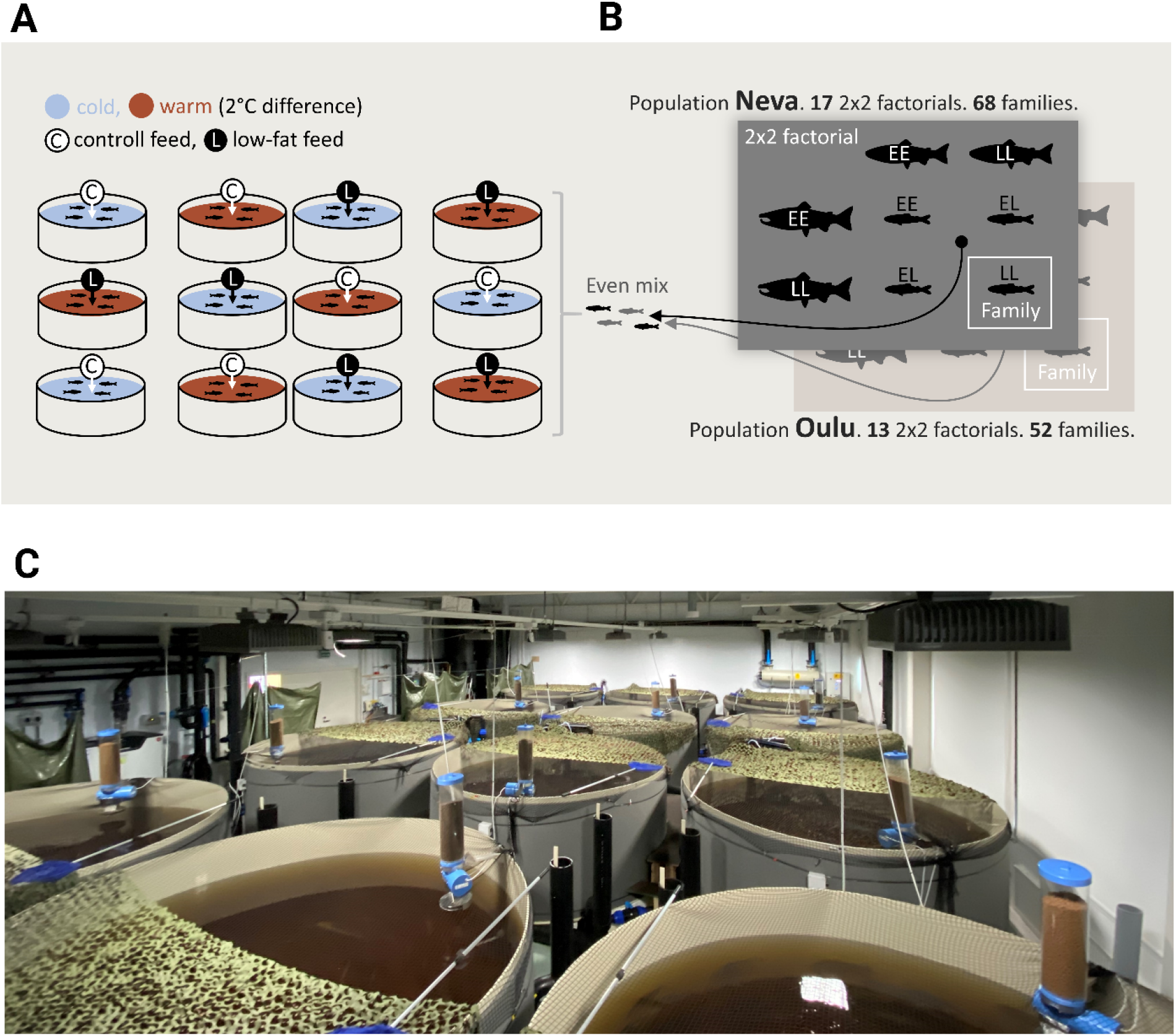
Design. Crossing, rearing, and experimental design. A) Shows the experimental tanks, their temperature- and feed treatments, as well as their relative positions in the room. B) Shows the crossing design, where unrelated parents from either the Neva or Oulu population were crossed in a series of 2 × 2 factorials (one *vgll3*EE* male and female and one *vgll3*LL* male and female) so that each 2 × 2 factorial yielded four families, one of each of the four reciprocal *vgll3* genotypes (EE, EL, LE or LL), i.e., all offspring within a family had the same *vgll3* genotype. Only individuals from the same population were crossed together. Roughly equal numbers of individuals from each family and population were placed into the 12 experimental tanks. C) A photograph of the experimental tanks taken in March 2020, showing tanks, feeders, and camouflage nets covering the tanks. The system relies on a constant flow of water drawn directly from the local lake Pääjärvi, which is where the water’s brown colour comes from.

**Table S1.1.** Feed: Usage overview of different pellet sizes. The table shows the date of first use of each pellet size for the different temperature- and feeding treatments. Pellet sizes were matched to the growth of the fish in the respective treatments. For future reference, this table contains dates used for the entire experiment; dates with asterisks are not included in this particular study. Feeds with end use “…” were used until the end of the experiment, and dates marked * are outside the time-scope of this paper.

| Feed | Temp | Size (mm) | Start use | End use |
| --- | --- | --- | --- | --- |
| low fat | warm | 1,2 | 10.07.2019 | *16.08.2020 |
| low fat | warm | 1,7 | 10.07.2019 | *20.07.2020 |
| low fat | warm | 2,5 | 16.08.2019 | *16.08.2020 |
| low fat | warm | 3,5 | 01.09.2019 | *15.05.2021 |
| low fat | warm | 5 | 07.10.2019 | ... |
| low fat | warm | 7 | *01.05.2020 | ... |
| low fat | cold | 1,2 | 17.07.2019 | *20.12.2020 |
| low fat | cold | 1,7 | 17.07.2019 | *04.04.2021 |
| low fat | cold | 2,5 | 16.08.2019 | *04.04.2021 |
| low fat | cold | 3,5 | 09.09.2019 | *04.04.2021 |
| low fat | cold | 5 | 07.10.2019 | *04.04.2021 |
| low fat | cold | 7 | *25.05.2020 | *04.04.2021 |
| controll | warm | 0,8 | 23.02.2018 | 10.07.2019 |
| Controll | warm | 1,2 | 29.05.2019 | *16.08.2020 |
| Controll | warm | 1,7 | 29.05.2019 | *20.07.2020 |
| Controll | warm | 2,5 | 16.08.2019 | *08.11.2020 |
| Controll | warm | 3,5 | 01.09.2019 | *15.05.2021 |
| Controll | warm | 5 | 07.10.2019 | ... |
| Controll | warm | 7 | *01.05.2020 | ... |
| controll | cold | 0,8 | 10.03.2018 | 10.07.2019 |
| Controll | cold | 1,2 | 29.05.2019 | *20.12.2020 |
| Controll | cold | 1,7 | 29.05.2019 | *04.04.2021 |
| Controll | cold | 2,5 | 16.08.2019 | *04.04.2021 |
| Controll | cold | 3,5 | 09.09.2019 | *04.04.2021 |
| Controll | cold | 5 | 07.10.2019 | *04.04.2021 |
| Controll | cold | 7 | *25.05.2020 | *04.04.2021 |

**Table S1.2.** Nutrients: Nutritional overview of different feed types and pellet sizes, as well as intended fish-size range for different pellet sizes.

| Size(mm) | Fish-size (g) | Type | % Moisture | % Ash | % Fat | % Protein | % Carbs | Energy (kj/g) |
| --- | --- | --- | --- | --- | --- | --- | --- | --- |
| 0,8 | 0.8-4 | control | 4,8 | 8,4 | 16,9 | 57,6 | 12,3 | 18,1 |
| 1,2 | 4-15 | control | 5,4 | 7,2 | 17,7 | 53 | 16,7 | 18,4 |
| 1,2 | 4-15 | low fat | 6,3 | 6,8 | 13,4 | 53,3 | 20,2 | 17,5 |
| 1,7 | 15-30 | control | 6,3 | 7,3 | 21,6 | 48,9 | 15,9 | 19,0 |
| 1,7 | 15-30 | low fat | 8,4 | 7,9 | 13,7 | 51,1 | 18,9 | 17,0 |
| 2,5 | 25-70 | control | 5,5 | 6,6 | 22,3 | 47 | 18,7 | 19,4 |
| 2,5 | 25-70 | low fat | 6,0 | 7,3 | 12,8 | 52,2 | 21,7 | 17,3 |
| 3,5 | 60-125 | control | 4,9 | 5,8 | 26,2 | 45,3 | 17,8 | 20,4 |
| 3,5 | 60-125 | low fat | 5,6 | 6,9 | 11,7 | 53,6 | 22,2 | 17,2 |
| 5 | 110-500 | control | 7,7 | 4,9 | 29,7 | 38,4 | 19,6 | 20,8 |
| 5 | 110-500 | low fat | 8,3 | 6,5 | 9,7 | 50,3 | 25,8 | 16,5 |
| 7 | 450-2300 | control | 5,9 | 4,3 | 33,7 | 35,8 | 20,0 | 22,0 |
| 7 | 450-2300 | low fat | 6,4 | 6,1 | 9,0 | 48,8 | 29,3 | 16,6 |

## SUPPLEMENTARY MATERIAL 2: Supplementary results

**Figure S2.1.**
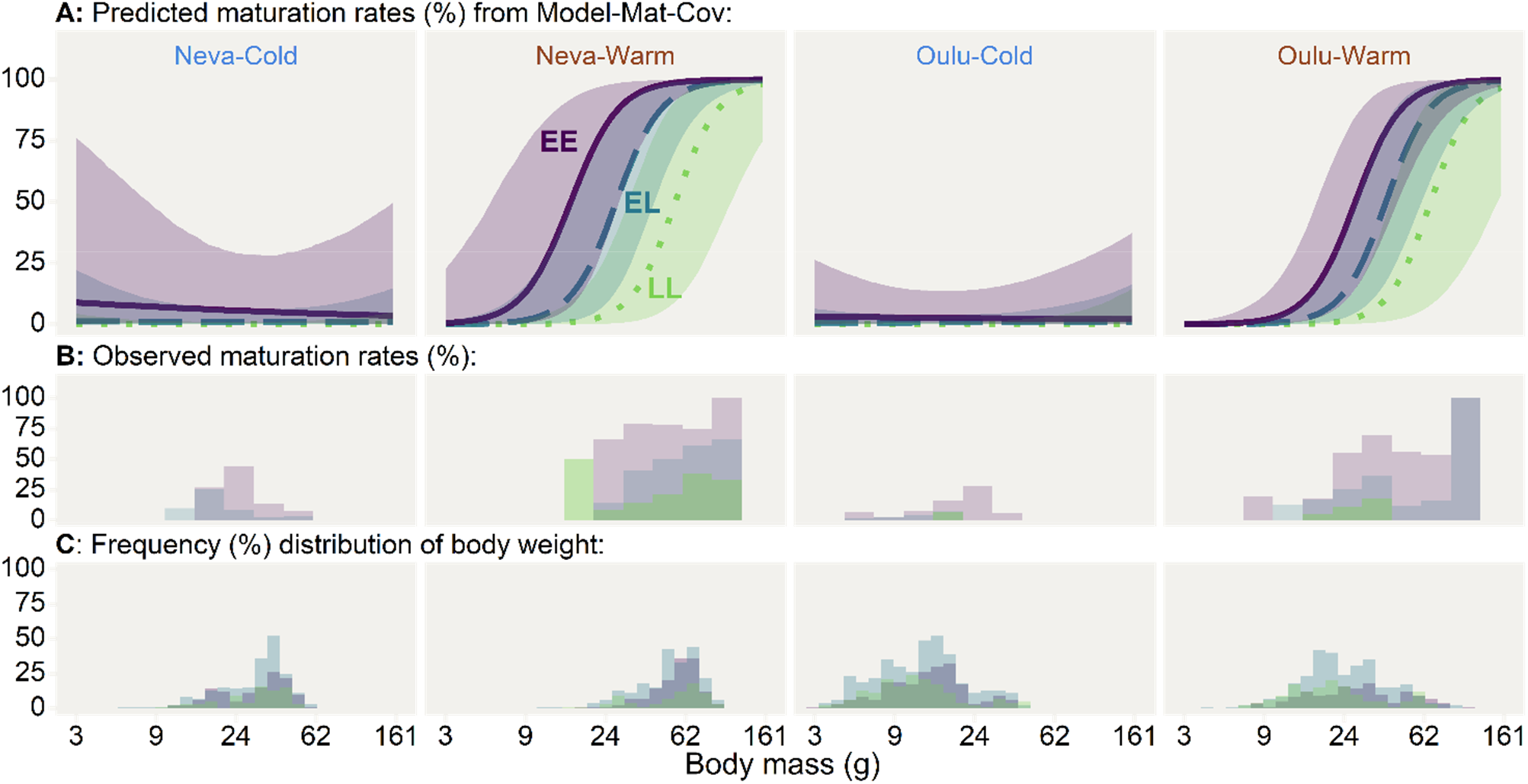
Maturation. Predicted maturation probabilities(A) and observed (B) maturation rates for three different *vgll3* genotypes (Purple-solid=EE, Blue-dashed=EL, Green-dotted=LL), two temperature treatments (Cold, Warm), and two populations of origin (Neva, Oulu), plotted against log-scaled body mass together with frequency distributions of body condition (C). x-axis labels are back-transformed from log scale to represent body mass in grams. Lines represent the mean predicted maturation probability for a given body mass, with shaded areas around the lines indicating the 95% credible interval for the predictions (overlap is not an indicator of significance). Predictions are based on the full model for maturation probability (Model-Mat-Cov, Figure 3-A, Table S2.4). For the model predictions, body condition here is to the mean of the whole study population and the migration-phenotype parameter is set to 0.5 (giving an estimate lying between migrant- and resident-phenotype individuals).

**Table S2.1.** CondObs. Observed body conditions for male Atlantic salmon in the summer of their second-year post-fertilization in different grouped combinations of *vgll3* genotype, temperature treatments and population of origin. Numbers after ± indicate SEM. Body condition is presented as percentage difference from predicted body mass (given body length). Numbers in parenthesis indicate the total number of fish in that group.

| Group | LL | EL | EE | Total |
| --- | --- | --- | --- | --- |
| Neva | 1.55%±0.54 (147) | 1.99%±0.40 (422) | 2.99%±0.48 (266) | 0.02%±0.00 (835) |
| Oulu | 0.14%±0.67 (310) | -0.59%±0.35 (673) | 1.34%±0.48 (352) | 0.00%±0.00 (1335) |
| Warm | 2.24%±0.84 (215) | 1.45%±0.41 (517) | 3.25%±0.56 (284) | 0.02%±0.00 (1016) |
| Cold | -0.87%±0.53 (242) | -0.53%±0.35 (578) | 1.03%±0.41 (334) | 0.00%±0.00 (1154) |
| Neva, warm | 0.51%±0.79 (73) | 0.10%±0.57 (217) | 0.67%±0.72 (140) | 0.00%±0.00 (430) |
| Neva, cold | 2.58%±0.73 (74) | 3.99%±0.52 (205) | 5.57%±0.54 (126) | 0.04%±0.00 (405) |
| Oulu, warm | 3.13%±1.20 (142) | 2.42%±0.56 (300) | 5.75%±0.82 (144) | 0.03%±0.00 (586) |
| Oulu, cold | -2.39%±0.66 (168) | -3.02%±0.40 (373) | -1.72%±0.48 (208) | -0.03%±0.00 (749) |
| Total | 0.59%±0.49 (457) | 0.40%±0.27 (1095) | 2.05%±0.34 (618) | 0.01%±0.00 (2170) |

**Table S2.2.** SmoltObs. Observed smolting rates for male Atlantic salmon at two years post-fertilization for different grouped combinations of *vgll3* genotype, temperature treatments and population of origin. Numbers in parenthesis indicate total number of fish in that group.

| Group | LL | EL | EE | Total |
| --- | --- | --- | --- | --- |
| Neva | 89.8% (147) | 88.6% (422) | 94% (266) | 90.5% (835) |
| Oulu | 55.8% (310) | 62.1% (673) | 67.9% (352) | 62.2% (1335) |
| Warm | 84.7% (215) | 90.9% (517) | 92.3% (284) | 90% (1016) |
| Cold | 50.8% (242) | 55.7% (578) | 68% (334) | 58.2% (1154) |
| Neva, warm | 94.5% (73) | 96.8% (217) | 98.6% (140) | 97% (430) |
| Neva, cold | 85.1% (74) | 80% (205) | 88.9% (126) | 83.7% (405) |
| Oulu, warm | 79.6% (142) | 86.7% (300) | 86.1% (144) | 84.8% (586) |
| Oulu, cold | 35.7% (168) | 42.4% (373) | 55.3% (208) | 44.5% (749) |
| Total | 66.7% (457) | 72.3% (1095) | 79.1% (618) | 73.1% (2170) |

**Table S2.3.** MassObs. Observed body mass (g) for male Atlantic salmon in the summer of their second-year post-fertilization in different grouped combinations of *vgll3* genotype, temperature treatments and population of origin. Numbers after ± indicate the SEM. Numbers in parenthesis indicate the total number of fish in that group

| Group | LL | EL | EE | Total |
| --- | --- | --- | --- | --- |
| Neva | 43.1±1.6 (147) | 42.7±0.9 (422) | 44.7±1.1 (266) | 43.4±0.6 (835) |
| Oulu | 18.6±0.8 (310) | 20.0±0.5 (673) | 22.0±0.8 (352) | 20.2±0.4 (1335) |
| Warm | 34.9±1.5 (215) | 38.2±0.9 (517) | 44.2±1.2 (284) | 39.2±0.7 (1016) |
| Cold | 19.0±0.8 (242) | 20.4±0.5 (578) | 21.1±0.7 (334) | 20.3±0.4 (1154) |
| Neva, warm | 55.1±2.3 (73) | 53.3±1.2 (217) | 56.5±1.2 (140) | 54.7±0.8 (430) |
| Neva, cold | 31.3±1.2 (74) | 31.5±0.8 (205) | 31.5±1.0 (126) | 31.5±0.5 (405) |
| Oulu, warm | 24.5±1.3 (142) | 27.2±0.8 (300) | 32.2±1.5 (144) | 27.8±0.7 (586) |
| Oulu, cold | 13.6±0.7 (168) | 14.3±0.4 (373) | 14.9±0.5 (208) | 14.3±0.3 (749) |
| Total | 26.5±0.9 (457) | 28.8±0.6 (1095) | 31.7±0.8 (618) | 29.1±0.4 (2170) |

**Figure S2.2.**
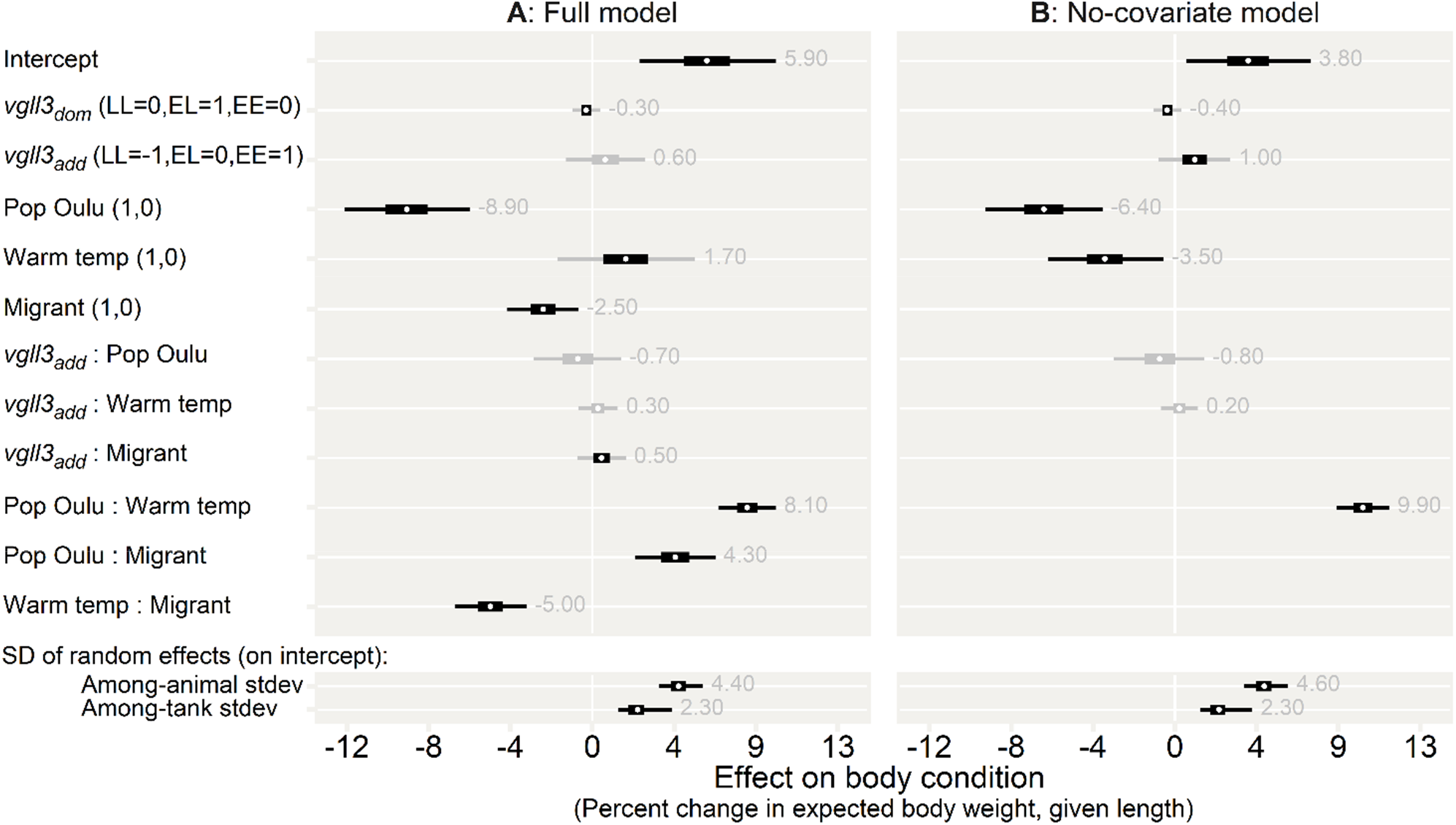
Model-Cond. Effect sizes (parameter estimates) for the two body condition models. A) Parameter estimates for the full model. B) A simplified model that only includes independent variables. Effects are transformed to show the percent change in expected body mass (given length), e.g., an effect size of 5 indicates that an individual is 5% heavier than what is expected for its mass. Thick and thin sections of bars indicate 97.5% and 50% credible intervals, respectively. As a visual aid, Intervals are coloured grey if they include 0. Grey numbers show the mean parameter estimate. Parentheses indicate the levels of the variables, and all variables are set to 0 for the intercept. The first level in parenthesis is the written level. The *vgll3dom* parameter indicates the degree of dominance displayed by either of the alleles. The lower section shows the standard deviation of the random effects, representing the degree of among-tank variation and among-animal variation (additive genetic standard deviation). The full model summaries can be found in Table S2.6 and S2.7.

**Figure S2.3.**
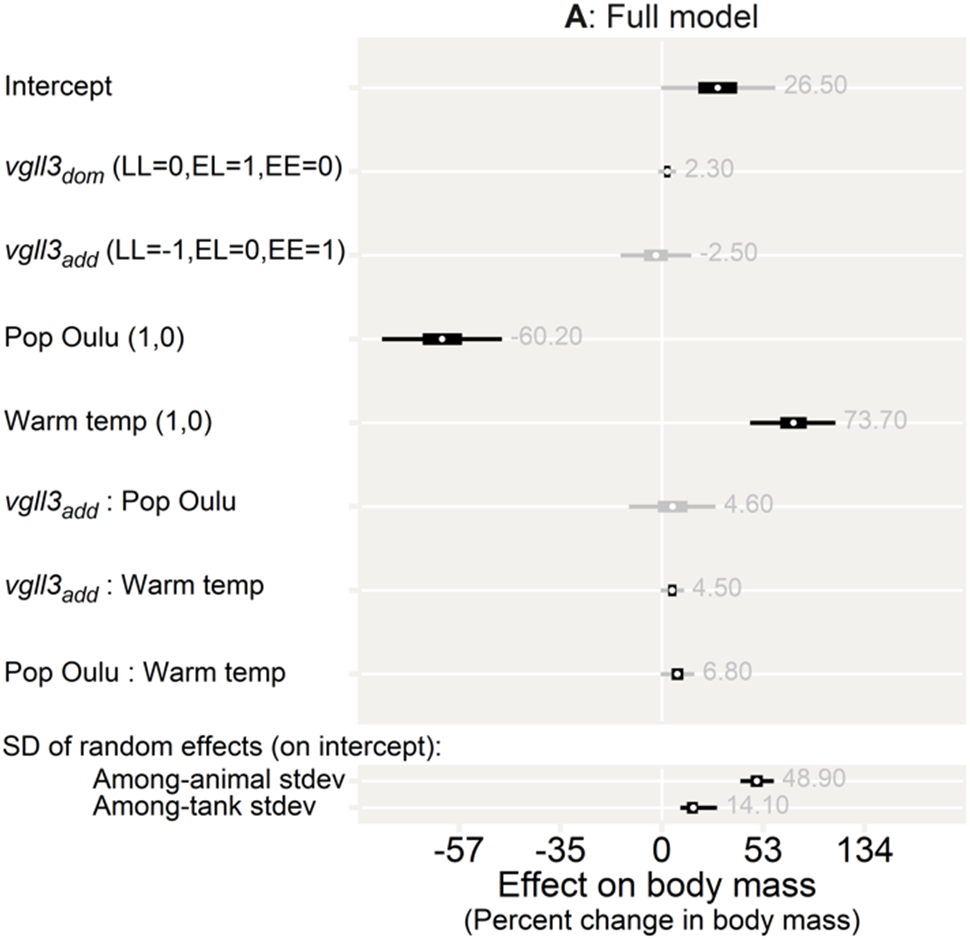
Model-Mass. Effect sizes (parameter estimates) for the body mass model. Effects are transformed to show the percent change in body mass. Thick and thin sections of bars indicate 97.5% and 50% credible intervals, respectively. As a visual aid, intervals are coloured grey if they include 0. Grey numbers show the mean parameter estimate. Parentheses indicate the levels of the variables, and all variables are set to 0 for the intercept. The first level in parenthesis is the written level. The *vgll3dom* parameter indicates the degree of dominance displayed by either of the alleles. The lower section shows the standard deviation of the random effects, representing the degree of among-tank variation and among-animal variation (additive genetic standard deviation). The full model summary can be found in Table S2.8

**Figure S2.4.**
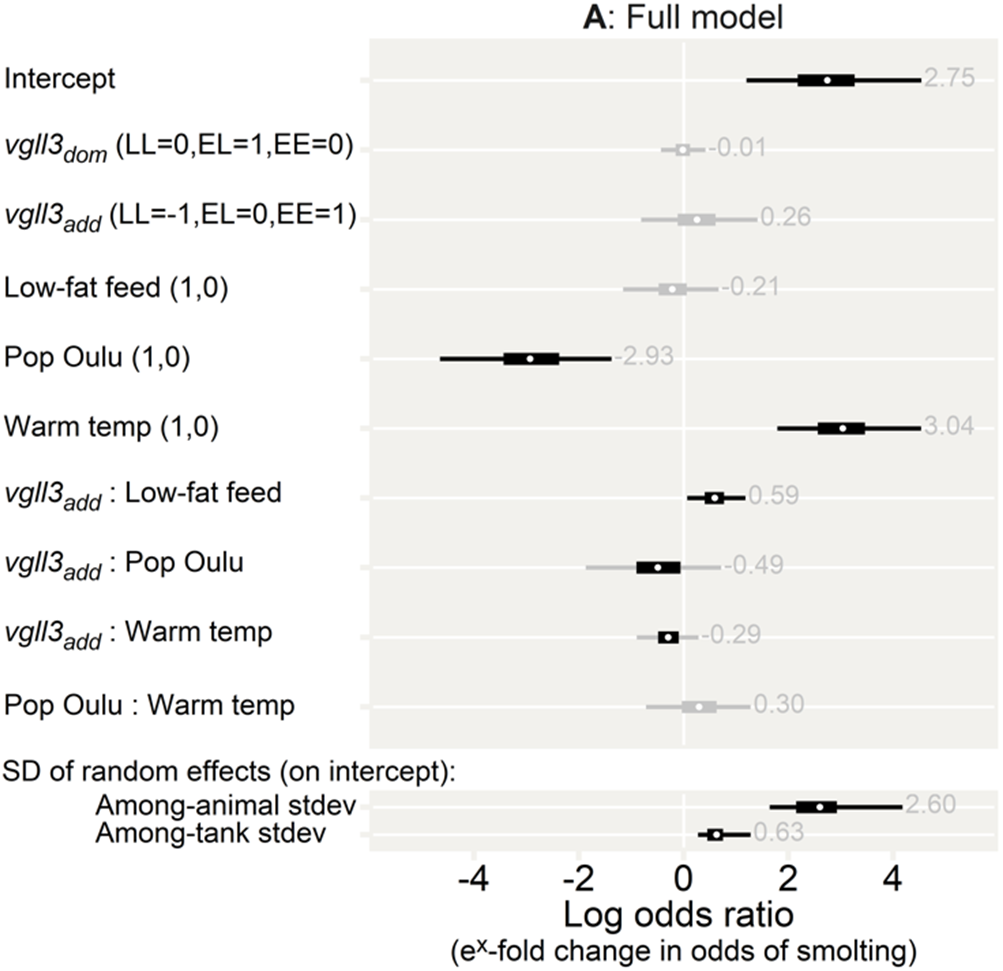
Model-MigPheno. Effect sizes (parameter estimates) for the migration phenotype model. Effects are shown as log odds ratios, thus indicating relative change in odds of transitioning from the resident (parr) to migrant (smolt) phenotype. Thick and thin sections of bars indicate 97.5% and 50% credible intervals, respectively. As a visual aid, intervals are coloured grey if they include 0. Grey numbers show the mean parameter estimate. Parentheses indicate the levels of the variables, and all variables are set to 0 for the intercept. The first level in parenthesis is the written level. The *vgll3dom* parameter indicates the degree of dominance displayed by either of the alleles. The lower section shows the standard deviation of the random effects, representing the degree of among-tank variation and among-animal variation (additive genetic standard deviation). The full model summary can be found in Table S2.9.

**Table S2.4.**
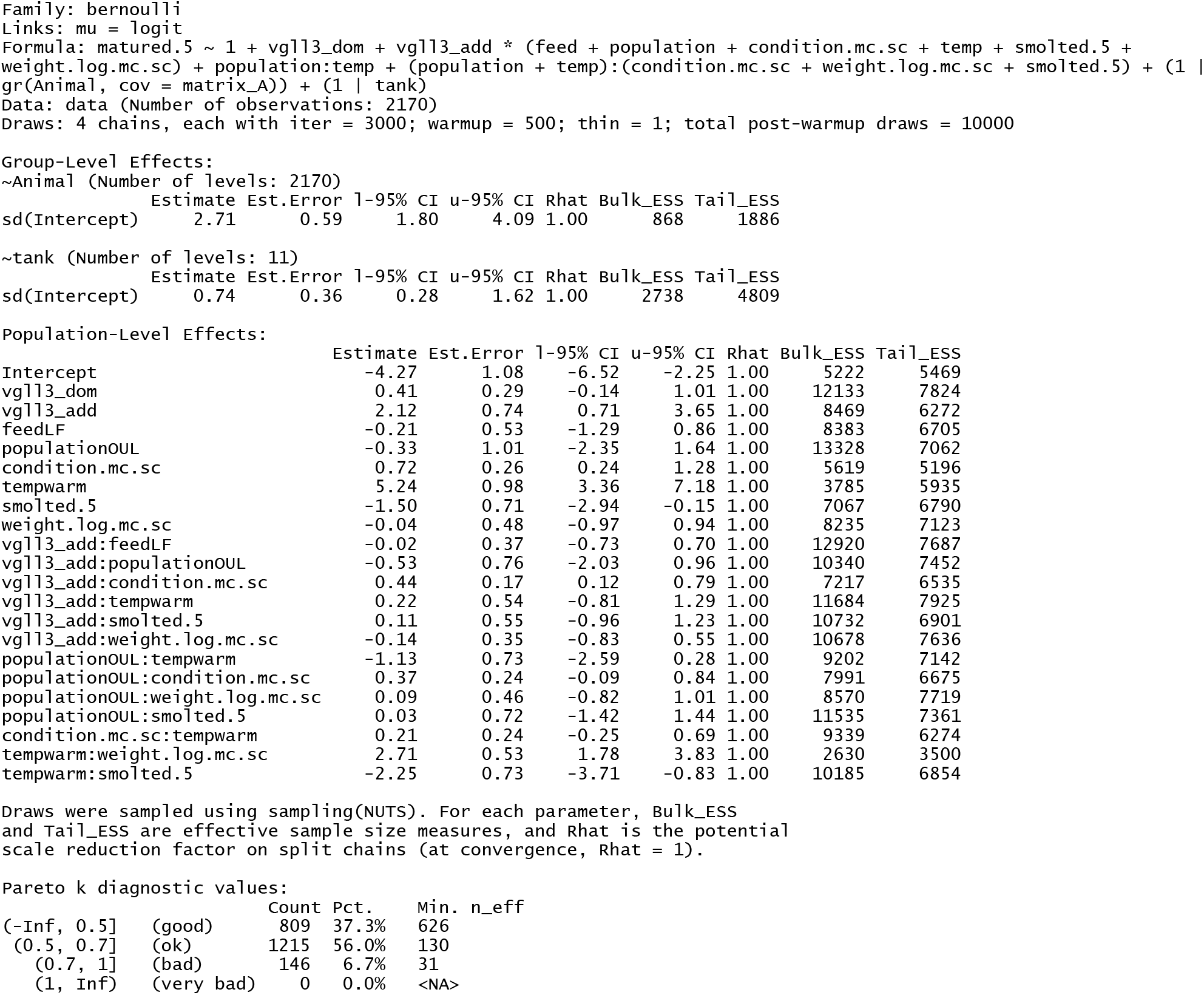
Summary-Model-Mat-Cov. Model summary output from *brms*, as well as pareto k diagnostic values from *loo* for the full maturation model (Model-Mat-Cov)

**Table S2.5.**
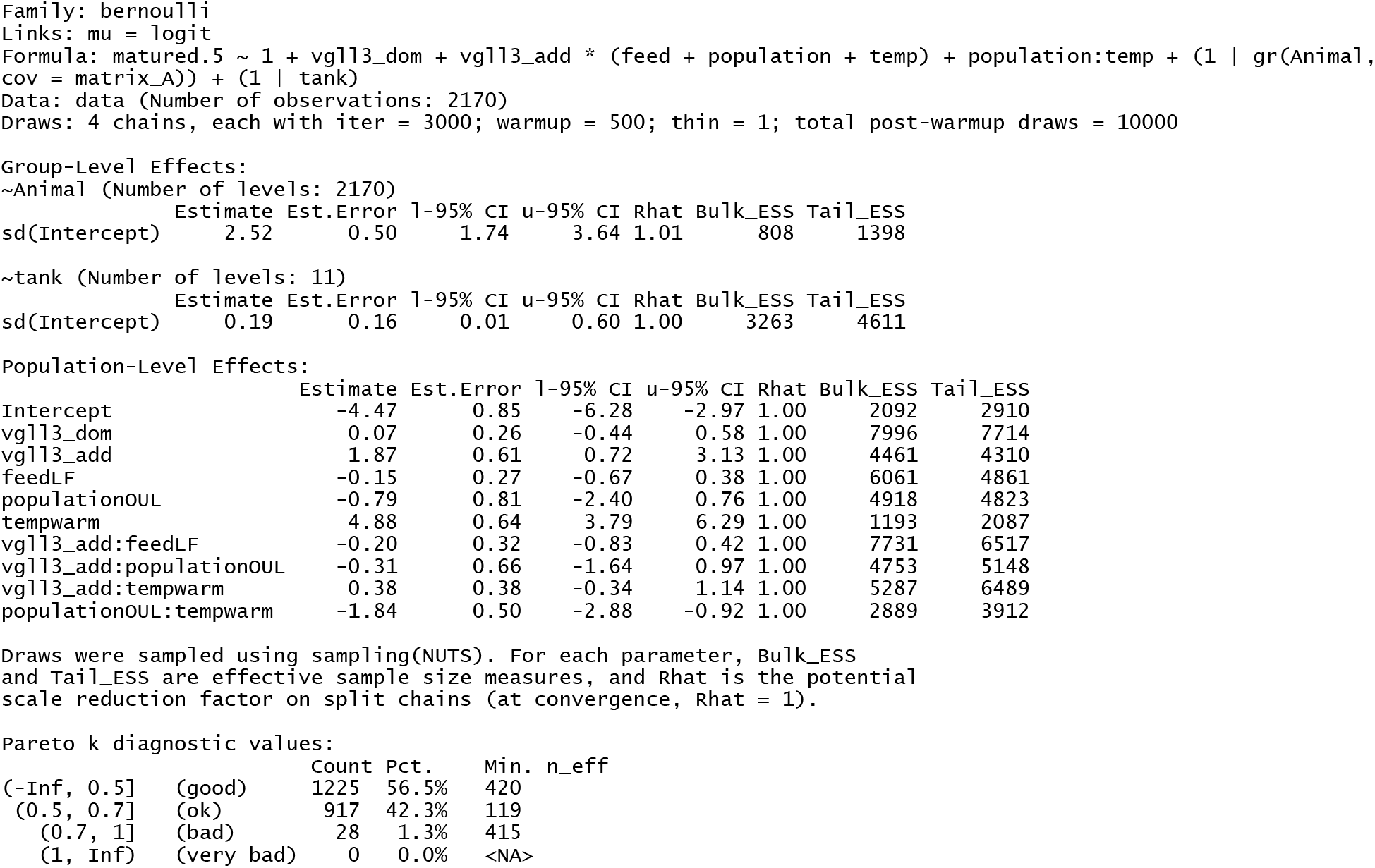
Summary-Model-Mat-Nocov. Model summary output from *brms*, as well as pareto k diagnostic values from *loo* for the no-covariate maturation model (Model-Mat-Nocov)

**Table S2.6.**
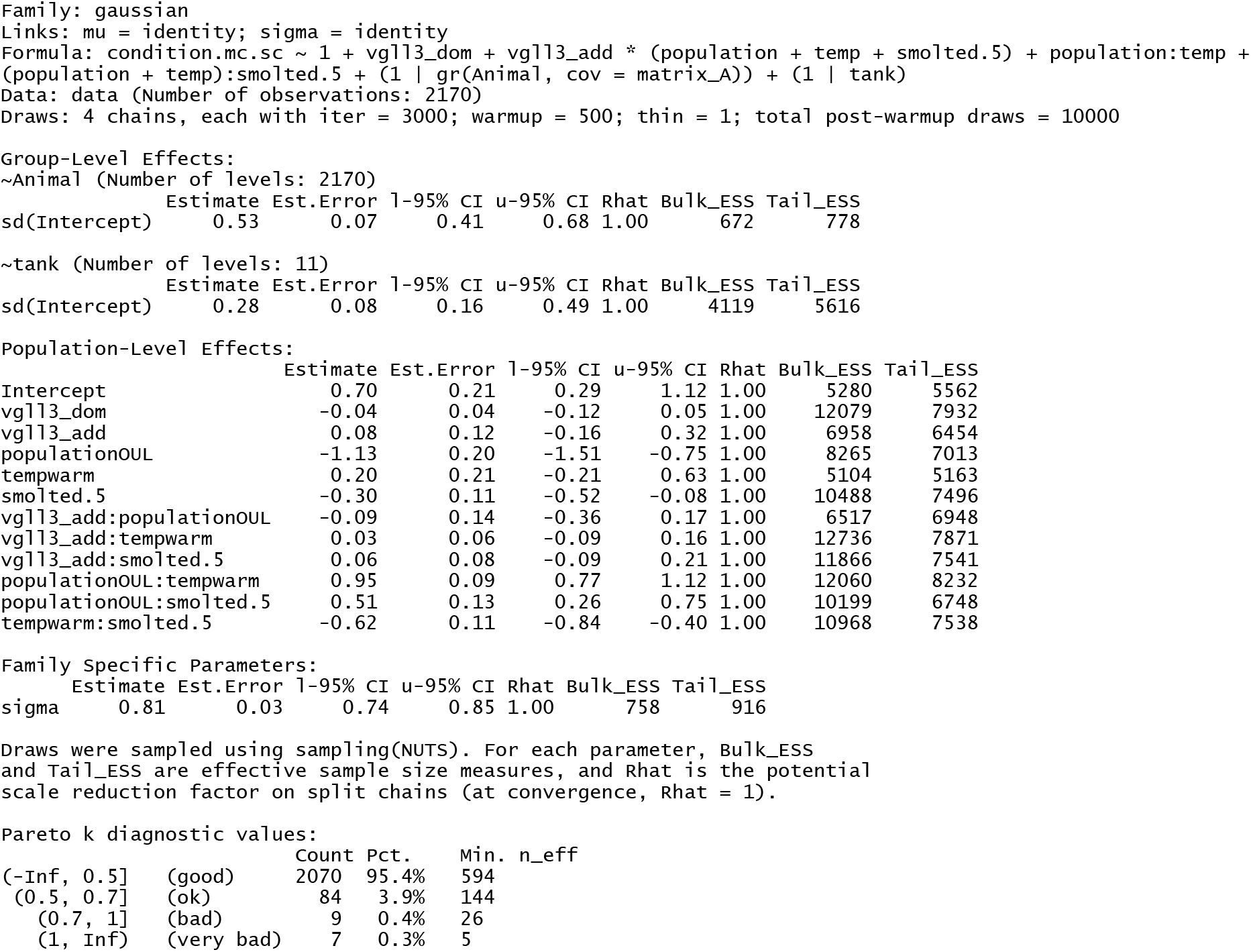
Summary-Model-Cond-Cov. Model summary output from *brms*, as well as pareto k diagnostic values from *loo* for the full body condition model (Model-Cond-Cov)

**Table S2.7.**
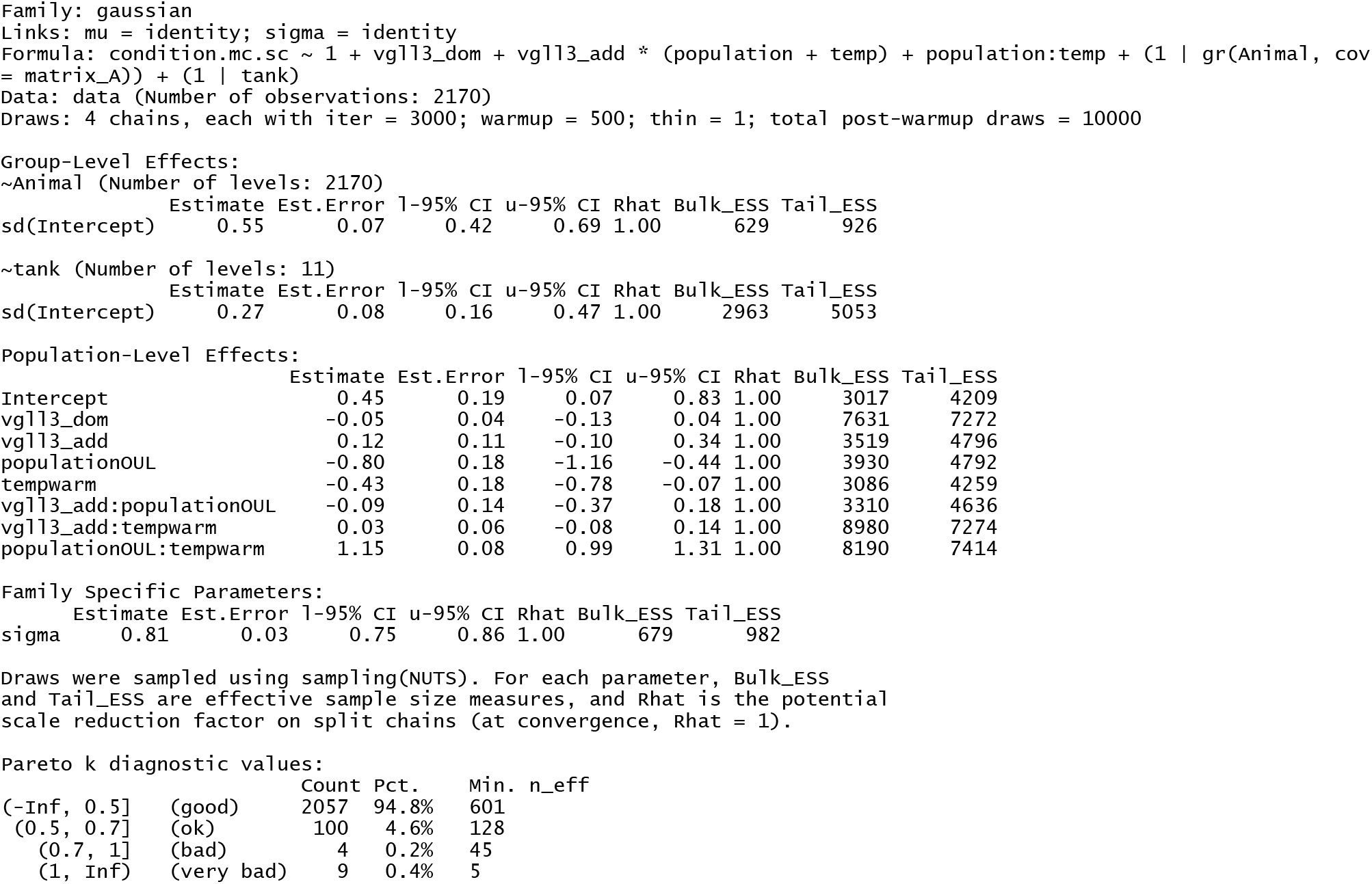
Summary-Model-Cond-Nocov. Model summary output from *brms*, as well as pareto k diagnostic values from *loo* for the no-covariate body condition model (Model-Cond-Nocov)

**Table S2.8.**
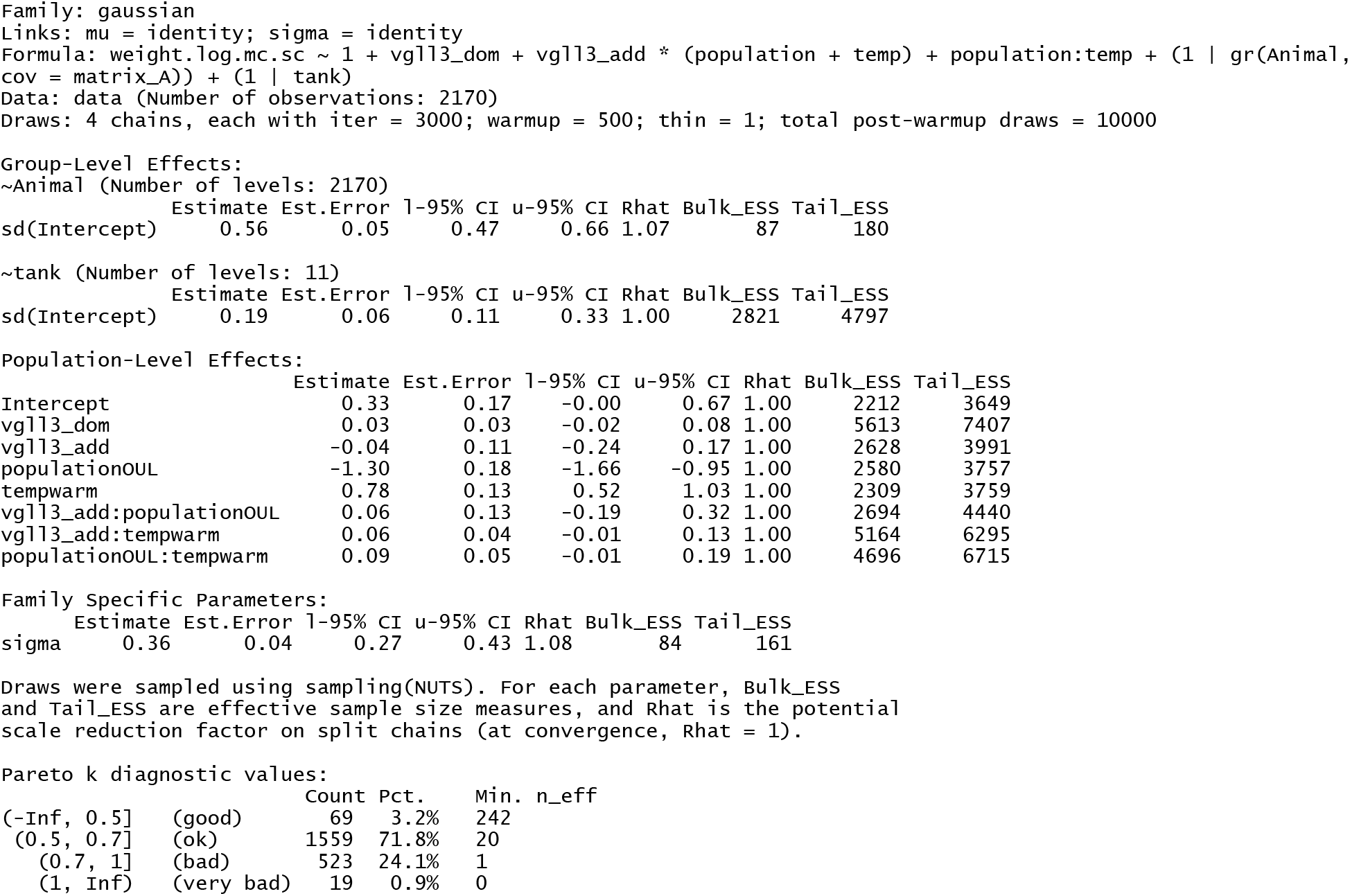
Summary-Model-Mass. Model summary output from *brms*, as well as pareto k diagnostic values from *loo* for the body mass model (Model-Mass)

**Table S2.9.**
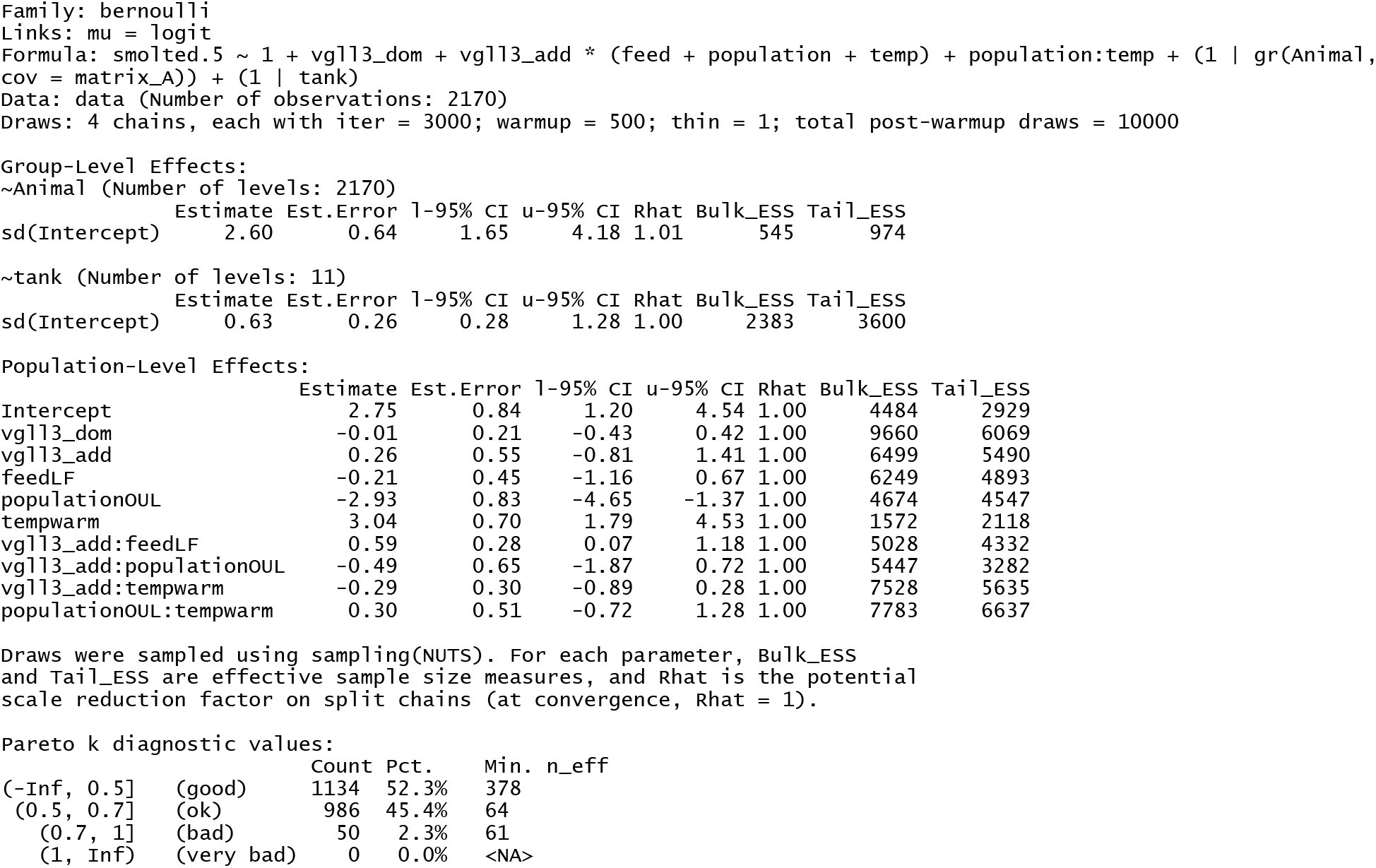
Summary-Model-MigPheno. Model summary output from *brms*, as well as pareto k diagnostic values from *loo* for the no-covariate body condition model (Model-MigPheno)

**Table S2.10.** Observed maturation of different combinations of maturation- and migration phenotypes (migrant=smolt, resident=parr) for male Atlantic salmon in the winter of their third year (early 2020) post-fertilization, for combinations of temperature treatment and population of origin.

| Group | n | Matured-migrant | Matured-Resident | Immature-Migrant | Immature-Resident |
| --- | --- | --- | --- | --- | --- |
| Neva, Cold | 405 | 6.9% | 2.7% | 76.8% | 13.6% |
| Neva, Warm | 430 | 52.3% | 1.9% | 44.7% | 1.2% |
| Oulu, Cold | 749 | 1.9% | 3.2% | 42.6% | 52.3% |
| Oulu, Warm | 586 | 17.7% | 6.0% | 67.1% | 9.2% |
| Neva | 835 | 30.3% | 2.3% | 60.2% | 7.2% |
| Oulu | 1335 | 8.8% | 4.4% | 53.3% | 33.4% |
| Cold | 1154 | 3.6% | 3.0% | 54.6% | 38.7% |
| Warm | 1016 | 32.4% | 4.2% | 57.6% | 5.8% |
| Total | 2170 | 17.1% | 3.6% | 56.0% | 23.3% |

